# Phylosymbiosis in the making: Discordant developmental emergence of evolutionary signatures in the mammalian gut microbiome and host transcriptome

**DOI:** 10.1101/2025.10.13.682227

**Authors:** Elizabeth N. Rudzki, Eldin Jašarević, Kevin D. Kohl

**Affiliations:** Biological Sciences Department, Dietrich School of Arts and Sciences, University of Pittsburgh, Pittsburgh, PA, USA; Department of Obstetrics, Gynecology, and Reproductive Sciences, School of Medicine, University of Pittsburgh, Pittsburgh, PA, USA; Magee-Womens Research Institute, Pittsburgh, PA, USA; Department of Computational and Systems Biology, School of Medicine, University of Pittsburgh, Pittsburgh, PA, USA; Department of Epidemiology and Biostatistics, Michigan State University, East Lansing, MI, USA

## Abstract

Gut microbial communities often mirror host phylogenetic relationships, an eco-evolutionary pattern known as phylosymbiosis. While phylosymbiosis is well documented in adult mammals, the developmental timing and mechanisms underlying its emergence remain unknown. Here, we use four captive-bred species of *Peromyscus* mice, a powerful model for studying natural variation in animal biology, to investigate the assembly of the intestinal microbiota and the emergence of phylosymbiotic signatures from birth through weaning. We combine 16S rRNA microbiome profiling with whole-tissue transcriptomics across three developmental stages (newborn, pre-weaning, and weaning) to test when during development the microbiome and host transcriptome exhibit evolutionary signatures. While parental feces and maternal mammary skin harbored phylosymbiotic communities, the intestines of newborn pups did not. Phylosymbiosis only emerged in pups at the pre-weaning stage of development and was sustained through weaning. Gut microbial diversity increased with age across species, and developmental stage was a stronger driver of community composition than host species identity, with a 1.5-2x larger effect size. Strikingly, host gene expression profiles showed the opposite pattern: transcriptome-based dendrograms were congruent with host phylogeny only at birth, not at later stages. Gene co-expression modules that showed correlation with host phylogenetic distance at birth were mainly enriched for metabolic pathways, such as lysosome activity, fat and vitamin digestion and absorption, and glycan metabolism. This work provides the first evidence, to our knowledge, in vertebrates that phylosymbiosis is not a fixed property of host-microbiome systems, but an emergent feature with distinct developmental dynamics in both the microbiome and the host transcriptome. Our results begin to reveal the complex and dynamic nature by which mammalian host-microbiome relationships are established and maintained across both evolutionary and developmental and time scales.

## Introduction

Mammals share close evolutionary ties to the microbial communities inhabiting their gastrointestinal tracts [1]. Many animals, including mammals, exhibit strong patterns of phylosymbiosis, in which the composition and structure of gut microbial communities mirror evolutionary relationships of their hosts [2-4]. These phylosymbiotic patterns carry significant functional consequences for the host [2]. While phylosymbiosis has been well documented in adult mammals [2, 5, 6], the timing and mechanisms by which these patterns first emerge during early development are unclear. Understanding the forces that shape the gut microbiota during early host development is a key open question in the field [7, 8].

The assembly of the gut microbiota is governed by several ecological processes, including selection, dispersal, and diversification [9, 10]. Host traits can act as selective filters on the assembling microbial community, such as gut pH [11, 12], mucin composition [13], gut motility [14], and immune function [15-17]. Dispersal also significantly influences microbiota acquisition, with newborns acquiring microbes through horizontal [18-21] and vertical transmission [22-24]. However, the specific processes driving phylosymbiosis remain unclear, as most studies have focused on assessing microbial dynamics at a single time point in adulthood.

Untangling the mechanisms of community assembly and phylosymbiosis is challenging because host-microbiome interactions are both complex and bidirectional. Gut microbial diversity increases during postnatal development [25, 26], and this diversification coincides with rising levels of bacterial metabolites such as short-chain fatty acids [27]. These microbial-derived metabolites influence host gene expression [28, 29], while hosts, in turn, regulate their microbiota through specific gene expression programs [30]. This reciprocal relationship indicates that examining either side of host-microbe crosstalk yields an incomplete understanding.

Here, we use four species of captively-bred *Peromyscus* mice to characterize how the intestinal microbiota develops and how phylosymbiotic signatures emerge over early life. *Peromyscus* mice are a well-established model for studying natural variation and the evolution of animal physiology, disease, and behavior [31-33]. Stocks of multiple captive-bred species are maintained at the Peromyscus Genetic Stock Center (PGSC), and these species differ in life-history traits such as developmental tempo and reproductive fitness (Section A in S1 Appendix). With multiple annotated *Peromyscus* genomes now available, this system provides a powerful opportunity to disentangle host genetic from environmental factors in shaping the microbiota. Specifically, we sampled the gut microbiome at several developmental time points and tracked how microbial community composition changes across species. We also performed transcriptomic sequencing of intestinal tissue and tested for evolutionary congruence of host gene expression across early life. Additionally, we investigated for timepoint-specific gene co-expression modules whose expression patterns correlated with host phylogenetic distance.

## Results

### Gut microbial diversity and composition shift dramatically across early development

We first evaluated whether alpha diversity of gut microbial communities increases as all four *Peromyscus* species age. We found that bacterial richness and evenness differed significantly across developmental timepoints in *P. californicus* (*CA*), *P. maniculatus* (*MA*), and *P. polionotus* (*PO*) (Kruskal-Wallis: Fig 1A, P = 0.0001; 1C, P = 0.0037; 1D, P = 0.0002; 1E, P < 0.0001; 1G, P = 0.0017; 1H, P = 0.0001). Pups of *CA* and *PO* showed a significant increasing monotonic trend with age for ASVs and Shannon diversity (Jonckheere-Terpstra: All P < 0.0046), however *MA* did not (all P > 0.091). Faith’s phylogenetic diversity and evenness showed similar age-related increases (Fig S1, Jonckheere-Terpstra: All P > 0.0048) for *CA* and *PO* pups, while only evenness was monotonically related to age for *MA* (Jonckheere-Terpstra: Faith’s PD P = 0.62, Evenness P = 0.032). Only Shannon diversity differed significantly by developmental timepoint for *P. leucopus* (*LE*) (Fig 1B, P = 0.064; 1F, P = 0.019, by Kruskal-Wallis), although all alpha diversity metrics did show a significant increasing monotonic trend with age (Jonckheere-Terpstra all P < 0.023), consistent with general patterns seen across most mammals. Full statistics, including Dunn’s post-hoc comparisons, are in Tables D-H in S1 Appendix.

**Fig 1.**
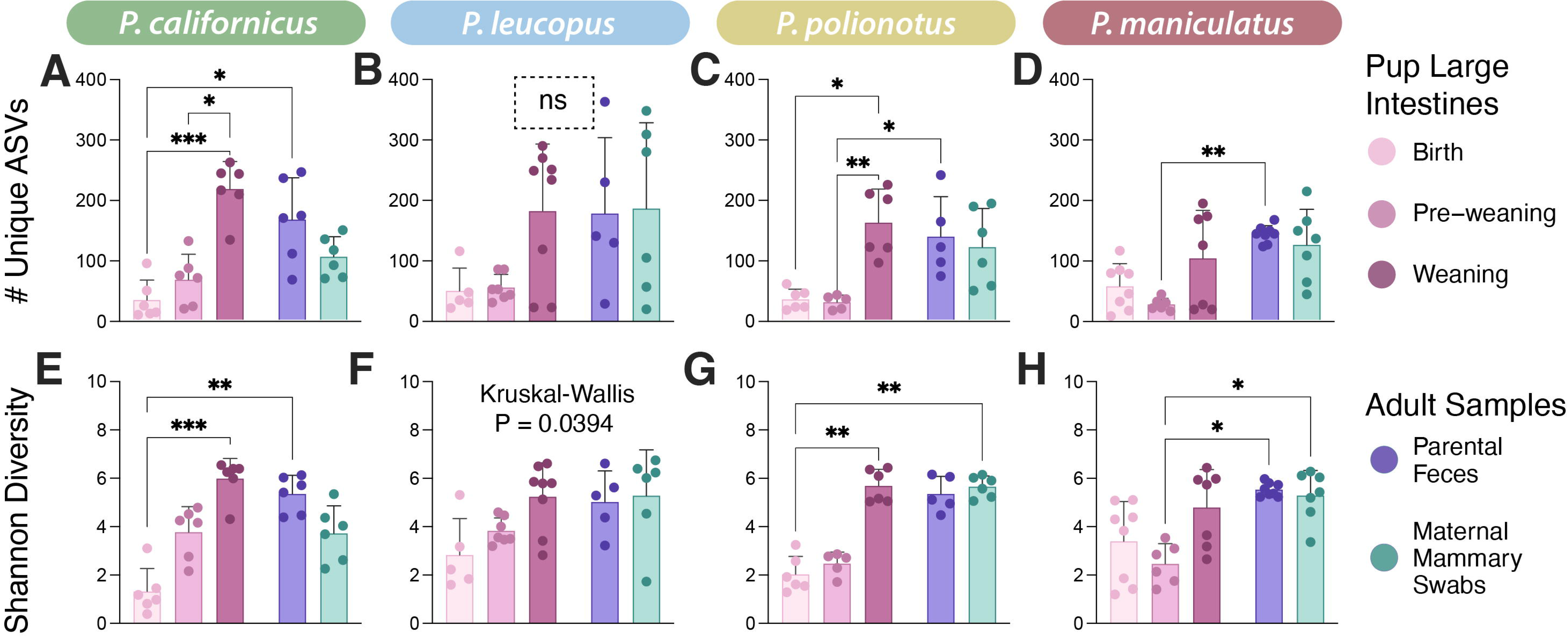
Alpha diversity metrics for the developmental comparison. Bacterial richness (# unique ASVs) present in each sample (repesented by points) for each of the four species **(A-D)**. Shannon diversity of the microbiota present in each sample for each of the four species **(E-H)**. Dunn’s multiple comparison statistical significance is represented with asterisks and brackets, when applicable.

Next, we evaluated beta diversity to confirm whether host species and/or developmental time point are major drivers of gut community composition. Both Jaccard (membership) and Bray-Curtis (structure) distances differed significantly across host species and sample types (adonis, P < 0.0001 for all; Table I in S1 Appendix). Sample type (encompassing developmental stage (pup and adult) and anatomical sampling site) had a 1.5-2x larger effect size (pseudo-F) than host species for both metrics. As pups aged, their gut microbiota became increasingly similar to the parental fecal microbiota (Fig S2). Within each species, gut community composition shifted significantly across pup developmental timepoints in both membership (adonis: *CA* P = 0.0268, *LE* P = 0.00332, *MA* P = 0.00027, *PO* P = 0.0075) and structure (*CA* P = 0.0045, *LE* P = 0.00061, *MA* P = 0.00131, *PO* P = 0.00099; Tables J and M in S1 Appendix). Cage ID did not significantly contribute to offspring microbial beta diversity.

Within-group variability (beta dispersion) also changed across development in certain species. *PO* and *MA* showed significant differences in dispersion by timepoint for both membership and structure (permadisp, P = 0.010 for all; Tables K-L and N-O in S1 Appendix), with newborn and weaning pups more variable than pre-weaning pups. Dispersion did not differ significantly across timepoints for *CA* or *LE* (membership: P = 0.39 and 0.94; structure: P = 0.18 and 0.25).

We then conducted differential abundance tests on dominant bacterial genera (>3% median relative abundance in any study group, Figs 2 and S3) to determine whether age-related gut microbial structure changes were driven by dominant members. Abundances of nearly all bacterial genera differed significantly between pup intestinal washes and adult fecal samples, at all pup timepoints (Kruskal-Wallis with B-H correction; adj. P < 0.0035), with the sole exception of *Aerococcus* (adj. P = 0.37). Several genera declined monotonically with age: *Streptococcus*, an unknown Corynebacteriaceae genus, an unknown Pasteurellaceae genus, *Escherichia-Shigella*, *Ligilactobacillus*, *Staphylococcus*, and *Rothia* (Jonckheere-Terpstra, B-H adj. P < 0.011). In contrast, genera showing monotonic increases in abundance included Muribaculaceae, two Lachnospiraceae genera (including *NK4A136* group), four Erysipelotrichaceae genera (including *Allobaculum* and *Ileibacterium*), two Lactobacillaceae genera, and *Helicobacter* (adj. P < 0.0091). Shifts in microbial structure across pup age were driven by differential abundances of dominant bacterial genera.

**Fig 2.**
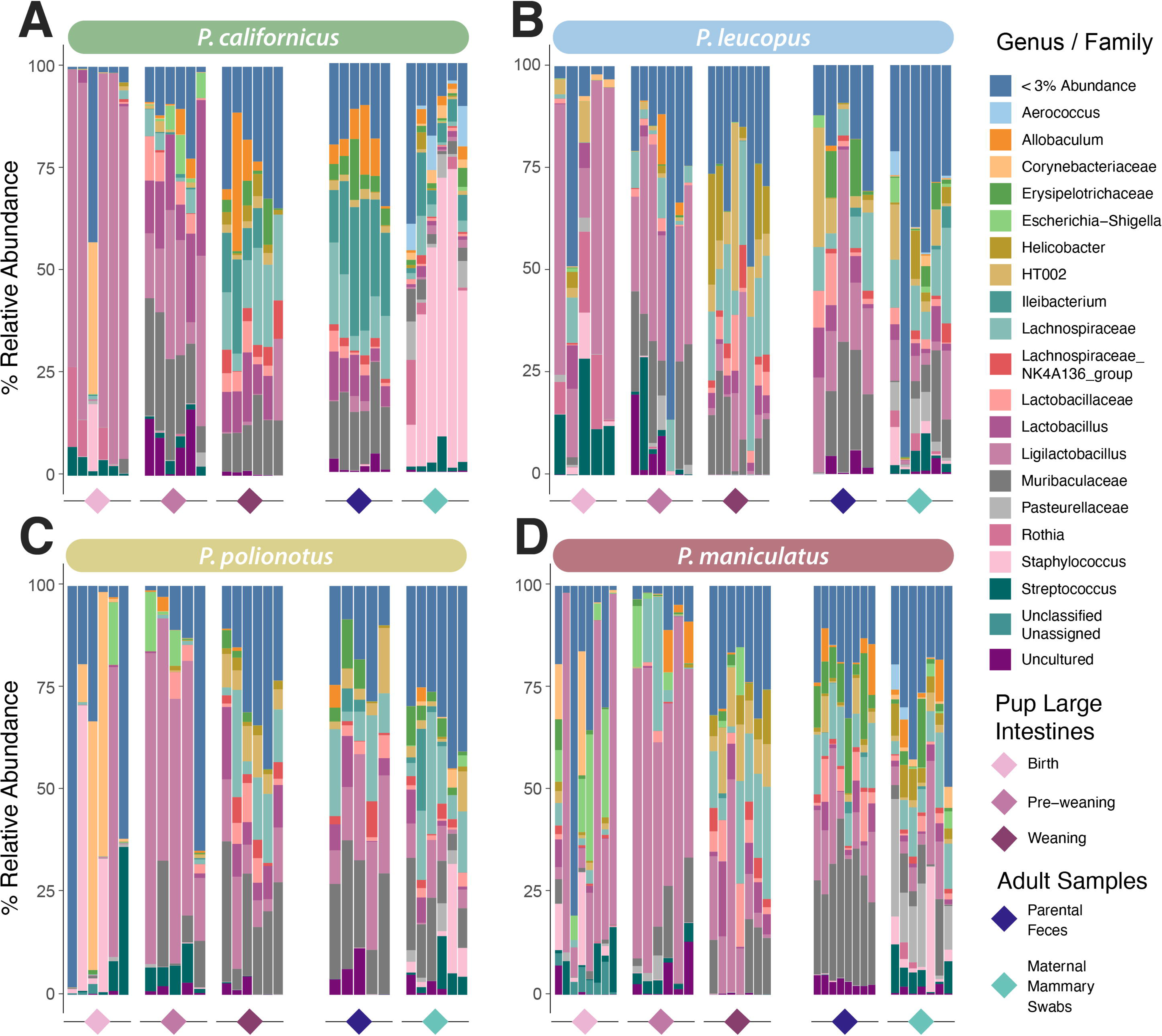
Taxa barplots showing the most abundant bacterial genera present in each sample. *P. californicus*. **(A)**, *P. leucopus* **(B)**, *P. polionotus* **(C)**, and *P. maniculatus* **(D)** developmental comparisons. The color of the diamond below each grouping of taxa barplots indicates the sample type (adult fecal, adult mammary skin, pup large intestinal wash from the three developmental timepoints).

Finally, we identified what bacterial genera differ significantly in abundance by host species or timepoint. *Allobaculum*, *Ileibacterium*, and *Rothia* differed significantly in relative abundance across host species (Kruskal-Wallis, B-H adj. P < 0.035, Figs 2 & S3). A Corynebacteriaceae genus and *Staphylococcus* were primarily detected on adult mammary skin (Corynebacteriaceae genus: 21/25, 1.3%; *Staphylococcus*: 23/25, 15%; for prevalence and mean relative abundance, respectively) and in newborn pup intestines (Corynebacteriaceae genus: 18/25, 8.9%; *Staphylococcus*: 18/25, 6.9%), then largely disappeared by later timepoints. The Corynebacteriaceae genus was present in only 1/27 weaning pups and 1/24 adult fecals at <0.04% abundances; *Staphylococcus* was present in 12/27 weaning pups and 14/24 adult fecals at <0.08% mean abundance). *Streptococcus* and the genus Pasteurellaceae were present in the guts of newborn pups (*Streptococcus*: 22/25, 6.7%; Pasteurellaceae: 14/25, 0.45%; for prevalence and mean relative abundance, respectively), pre-weaning pups (*Streptococcus*: 23/24, 3.7%; Pasteurellaceae: 14/24, 1.1%), and on adult mammary skin (*Streptococcus*: 25/25, 3.8%; Pasteurellaceae: 24/25, 6.0%), with minimal abundance in weaning pups (*Streptococcus*: 15/27, 0.034%; Pasteurellaceae: 5/27, 0.012%) and adult fecal samples (*Streptococcus*: 12/24, 0.024%; Pasteurellaceae: 0/24, 0%), suggesting a possible maternal transmission route. *Rothia* was largely restricted to newborn guts and adult mammary skin in *CA* (newborns: 4/6, 6.1%; adult mammary skin: 4/6, 3.8%; for prevalence and mean abundance, respectively) and *LE* (newborns: 5/5, 8.3%; adult mammary skin: 4/6, 1.5%). *MA* showed minimal presence/abundance in newborn guts (2/8, 0.12%) and adult mammary skin (2/7, 0.81%); and *PO* had only a single pup sample at the weaning timepoint where *Rothia* was detectable, at 0.02% abundance.

### Host gene expression is restructured across early development

We computed Euclidean distances from VST gene expression and visualized them using principal components analysis (Fig 3I-L) to determine whether host gene expression in the large intestine also changes markedly during early development. Developmental timepoint significantly explained gene expression variation in all four species (adonis: *CA* F = 9.78, P = 0.00076; *LE* F = 60.5, P < 0.0001; *MA* F = 13.9, P < 0.0001; *PO* F = 19.6, P < 0.0001; Table CC in S1 Appendix). A combined analysis of all pup samples showed that both developmental stage and host species identity are major drivers of the early-life intestinal transcriptome with similar effect sizes (Fig 5; developmental timepoint: F = 162, R² = 0.426; host species identity F = 123, R² = 0.483; Table BB in S1 Appendix). Beta dispersion differed by timepoint only in *PO* pups (permadisp, P = 0.040; Tables DD-EE in S1 Appendix); dispersion was stable across timepoints for the other three species (P > 0.25).

**Fig 3.**
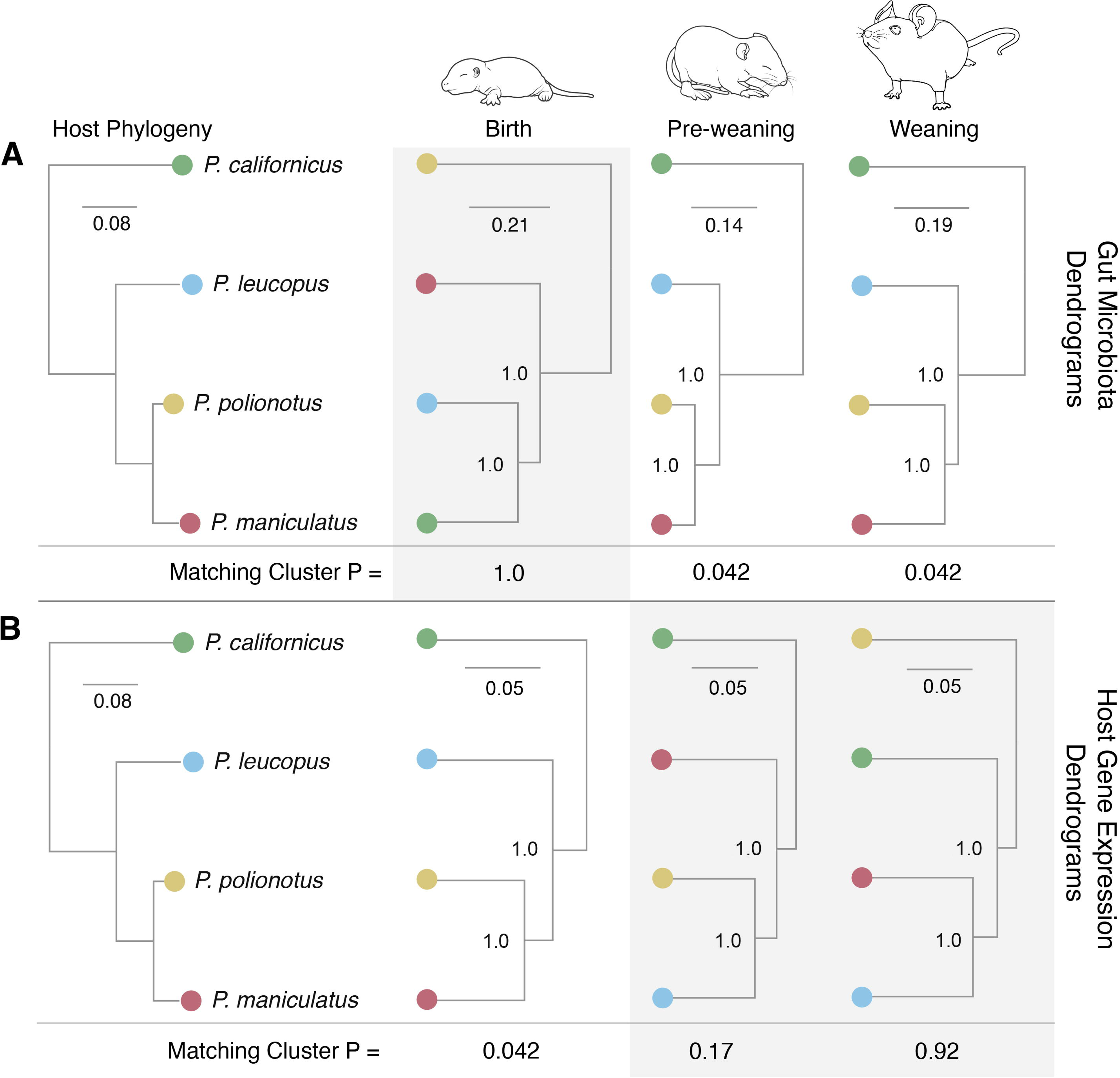
Beta diversity of microbiota and host transcriptomics for the developmental comparison. PcoA plots of beta diversity metrics for the developmental comparison of the bacterial microbiota dataset **(A-H)** for Jaccard similarity **(A-D)** and Bray-Curtis dissimilarity **(E-H)**. PCA plots of Euclidean distances for the developmental comparison of the host transcriptomics dataset **(I-L)**.

### Adult *Peromyscus* harbor phylosymbiotic microbial communities in fecal and mammary skin samples

We tested whether our adult *Peromyscus spp.* fecal and mammary skin samples show strong patterns of phylosymbiosis, where microbial community dendrograms are statistically congruent with host phylogeny. Consistent with prior reports [5], adult *Peromyscus* fecal communities showed significant patterns of phylosymbiosis (Matching Cluster (“MC”) P = 0.042). We also detected significant phylosymbiosis in the maternal mammary skin microbiota (MC P = 0.042), thereby extending this pattern to a body site not previously examined in this system.

### Phylosymbiosis emerges in the microbiome after phylogenetic structure in the transcriptome is lost

We next tested the primary question of this study, namely, when in early life do the gut microbiome and the host transcriptome acquire or lose evolutionary signatures? Phylosymbiosis in the gut microbiome emerged at the pre-weaning timepoint and was sustained at weaning (Fig 4A; MC P = 0.042). In contrast, host gene expression dendrograms were congruent with host phylogeny only at birth (Fig 4B; MC P = 0.042), with no significant congruence at later timepoints. Thus, similarities in transcriptional profiles reflect the host phylogenetic tree only at birth, whereas gut microbiome structures begin to reflect it at the preweaning stage, revealing discordant timing between the two. When feature tables were collapsed using the mean-ceiling method, results were broadly consistent with the median-ceiling approach, except that the weaning timepoint for the gut microbiota was no longer statistically significant (Fig S4, P = 0.17).

**Fig 4.**
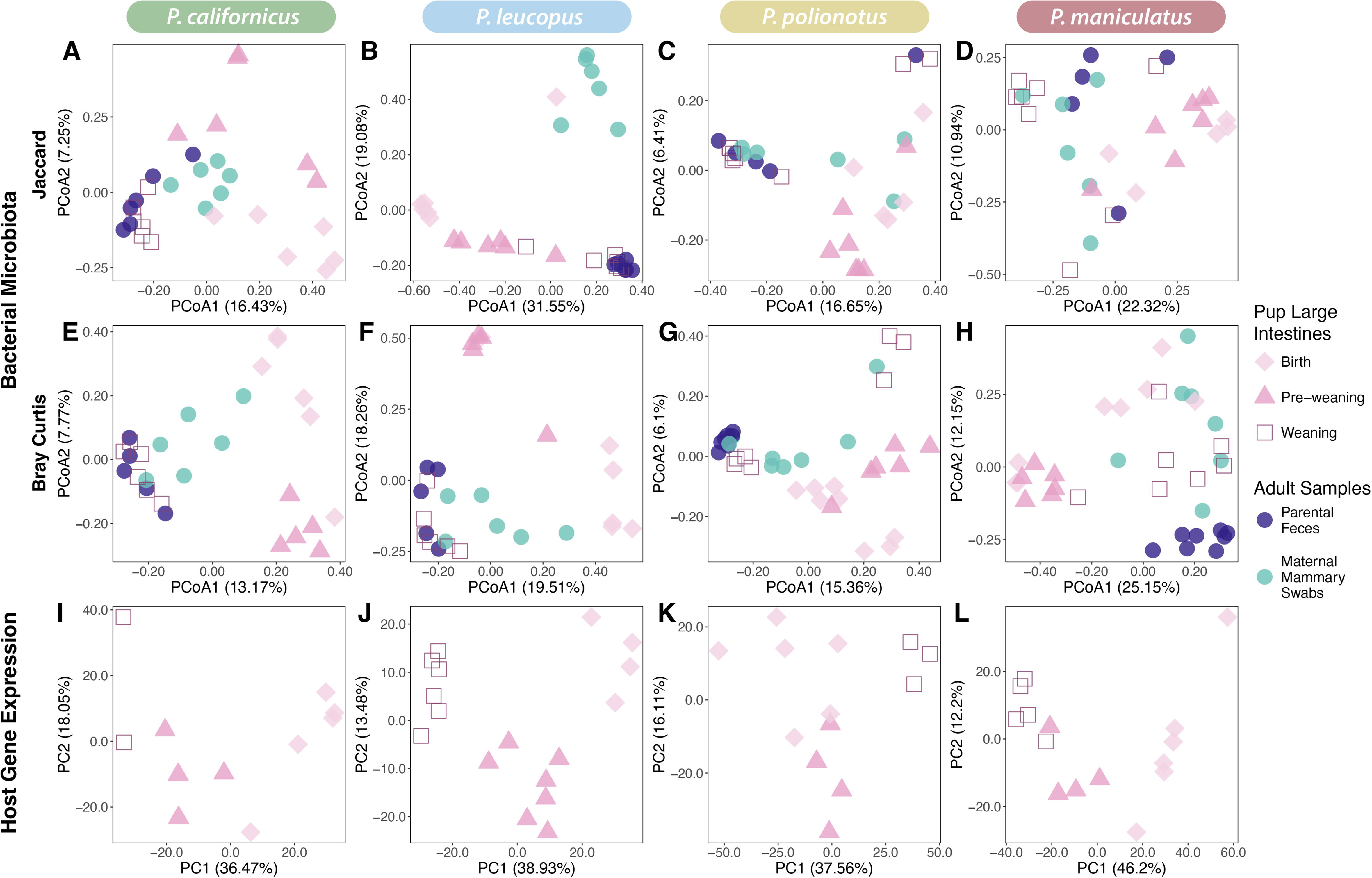
Dendrogram congruence of *Peromyscus* microbiota and gene expression profiles at different stages of host development. **(A)** Microbiota dendrograms and **(B)** host gene expression dendrograms were created from Bray-Curtis dissimilarity matrices (collapsed by host species by median-ceiling) and congruence compared to the host phylogenetic tree. Significance of the experimental dendrogram yielding an equal or better congruence to host phylogeny than that of 100,000 simulated random trees was calculated using matching cluster analysis. Congruent dendrograms with a significant normalized matching cluster score are highlighted with a white background. Rodent artwork was created and adapted from BioRender. Rudzki, E. (2026) https://BioRender.com/pvkioap

### Host species identity drives distinct microbiome and transcriptome profiles throughout development

We investigated the beta diversity of each host species at a given developmental timepoint to ascertain when *Peromyscus* pups begin harboring species-specific gut microbial communities. Host species identity significantly explained beta diversity for bacterial community membership (Fig 6A-D), bacterial community structure (Fig 6E-H), and host gene expression (Figs 5 and 6I-L; adonis, all P ≤ 0.0069; Tables FF, II, and LL in S1 Appendix). All four *Peromyscus* species harbored compositionally distinct gut microbial communities and intestinal gene expression profiles at each developmental time point (Fig 6). Taken together with the dendrogram congruence results (Fig 4), these data show that while the four species are compositionally distinct at every time point, only certain developmental windows yield profiles that mirror host phylogeny, and that this timing differs between the microbiome and the transcriptome.

**Fig 5.**
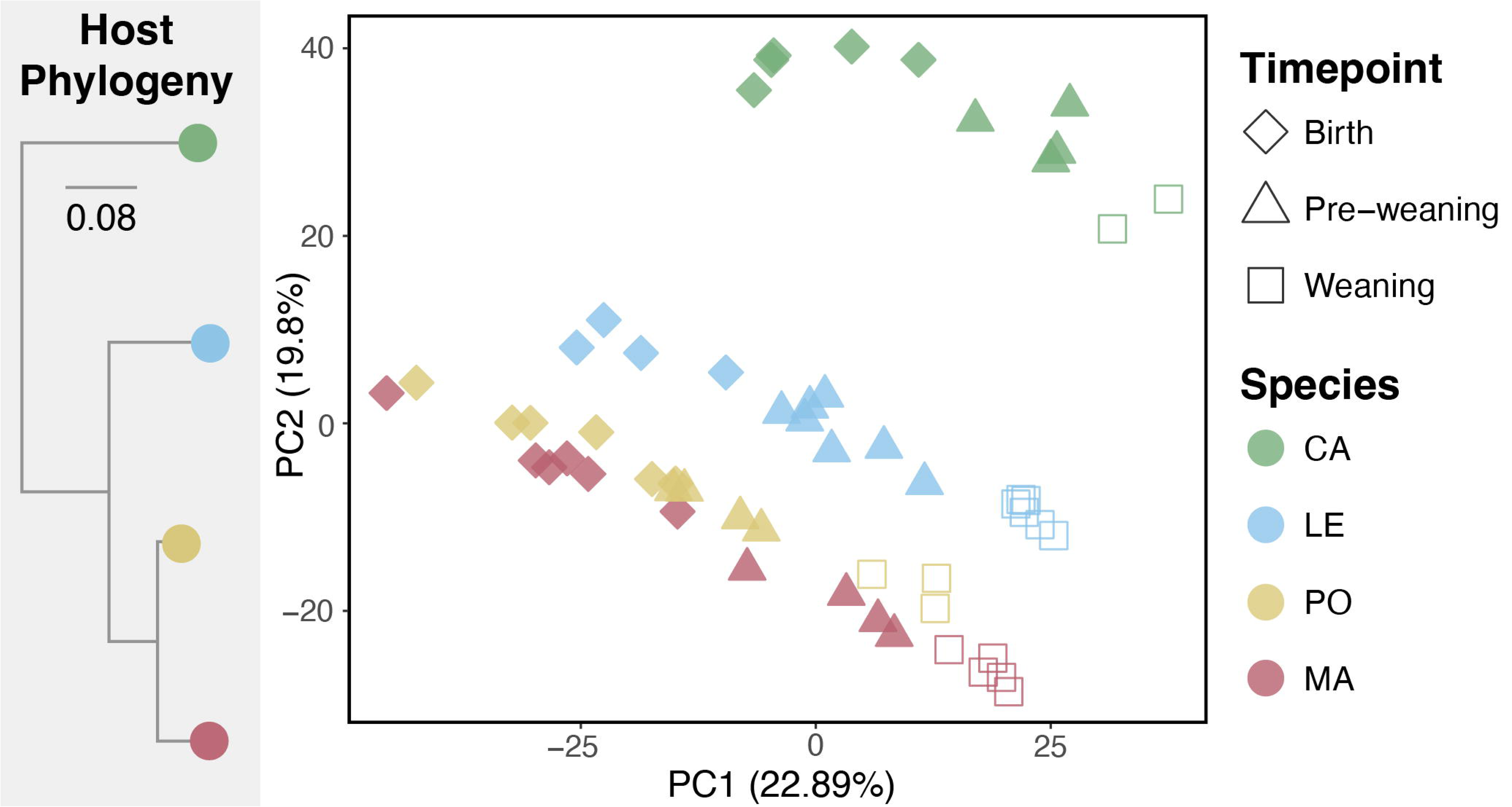
PCA plot of the entire host transcriptomics dataset.

**Fig 6.**
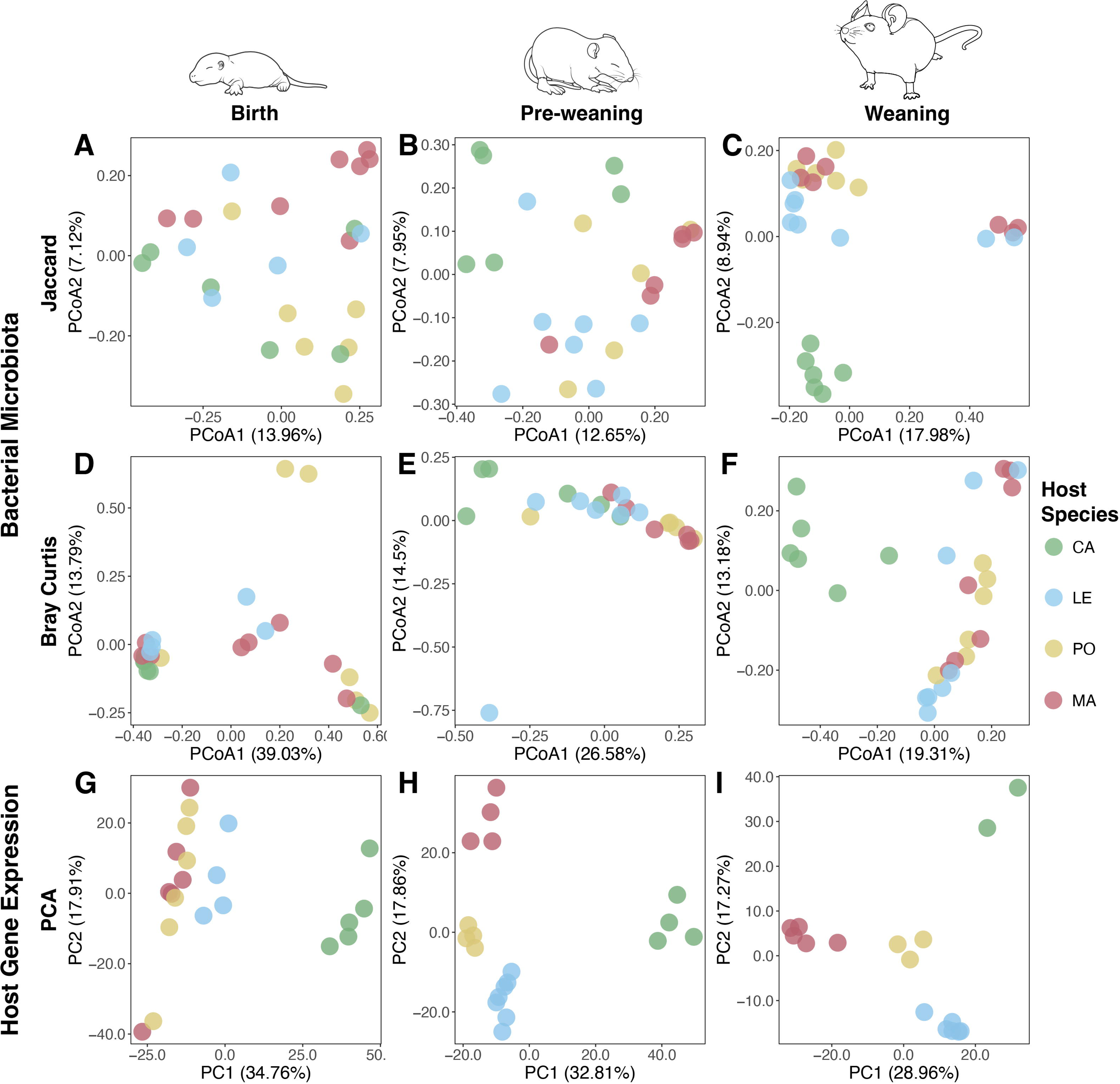
Ordination plots of microbiota and host transcriptomics for the host species comparison. PcoA plots of beta diversity metrics for the host species comparison of the bacterial microbiota dataset **(A-H)** for Jaccard similarity **(A-C)** and Bray-Curtis dissimilarity **(D-F)**. PCA plots of Euclidean distances for the host species comparison of the host transcriptomics dataset **(G-I)**.

Next, we performed differential gene expression analysis to evaluate broader trends in gene expression across our *Peromyscus* species as they age. Comparing gene expression across host species revealed numerous differentially expressed genes (DEGs) at early developmental time points, especially between more distantly related species pairs (e.g., *CA* versus others). As pups aged, DEG counts declined, and by weaning, the number of DEGs was broadly similar (mean 1,354 DEGs) across all species comparisons. We identified the top 50 most variable DEGs across all conditions (Fig 7). One cluster, including *Ceacam1, Tssk2, Oaz3,* and *Cryba4,* was more highly expressed in *LE*, *PO*, and *MA* than in *CA*. A second cluster, including *Xab2, Top1mt,* and *Tpt1*, was more highly expressed in *CA* than in the other three species, particularly at the weaning timepoint. These species-specific expression differences at weaning suggest that while overall DEG counts converge, specific functional gene networks remain divergent.

**Fig 7.**
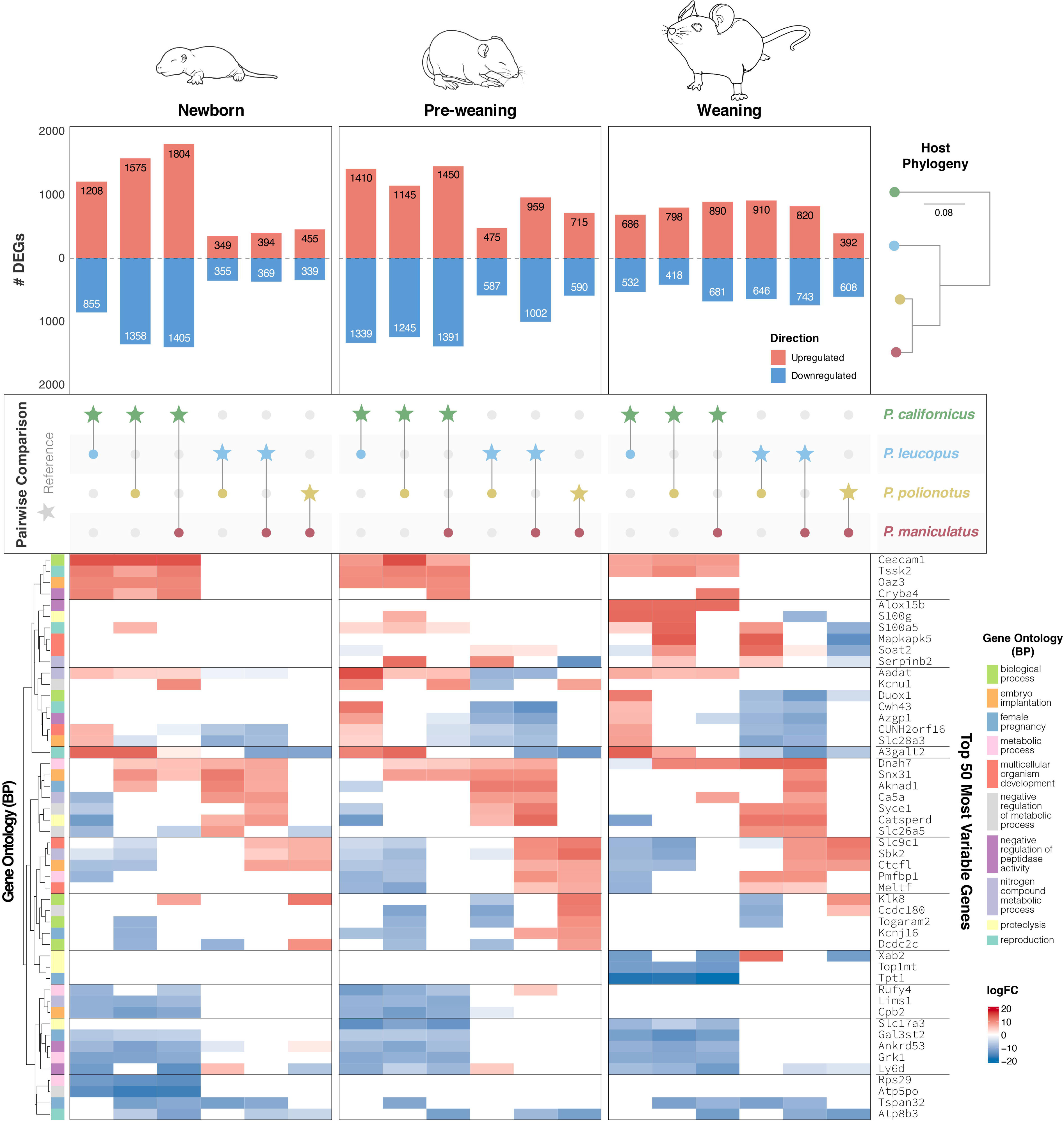
Genes differentially expressed between *Peromyscus* species at specific stages of host development. **Top**: Number of genes differentially expressed between each pairwise comparison. **Middle**: The two host species being compared, with the reference species notated as a star. **Bottom**: Heatmap of top 50 differentially expressed annotated genes with the highest variance across all conditions. Rodent artwork was created and adapted from BioRender. Rudzki, E. (2026) https://BioRender.com/pvkioap

### Newborn gut gene modules mirror host evolutionary relationships

We then used WGCNA to identify co-expression gene modules in the large intestinal transcriptome and tested whether any correlated with host phylogenetic distance. We identified 26 modules at the newborn timepoint and 30 at pre-weaning and assessed each using four correlational methods (Fig 8A). We detected more phylogenetically correlated modules at the newborn timepoint than at pre-weaning, consistent with the transcriptome-level congruence observed only at birth (Fig 4B). At the newborn timepoint, two modules met the most stringent criteria: “Orchid” (Mantel r = 0.962-0.964; all methods pre-FDR P = 0.0041-0.0833; bootstrapped weighted post-FDR P = 0.0291) and “Skyblue” (Mantel r = 0.972-0.973; all methods pre-FDR P = 0.0044-0.0833; bootstrapped weighted post-FDR adj. P = 0.0291). A third module, “Honeydew”, met the minimum criteria threshold (Mantel r = 0.916-0.962; weighted pre-FDR P = 0.0833; bootstrapped weighted pre-FDR P = 0.0273) (Fig 9B, first panel). At the pre-weaning timepoint, only one module, “Alabaster,” met the minimum threshold (Mantel r = 0.935-0.951; weighted pre-FDR P = 0.0833; weighted bootstrapped pre-FDR P = 0.0533) (Fig 8B, second panel). Full results for all gene modules are provided in the S2 Appendix.

**Fig 8.**
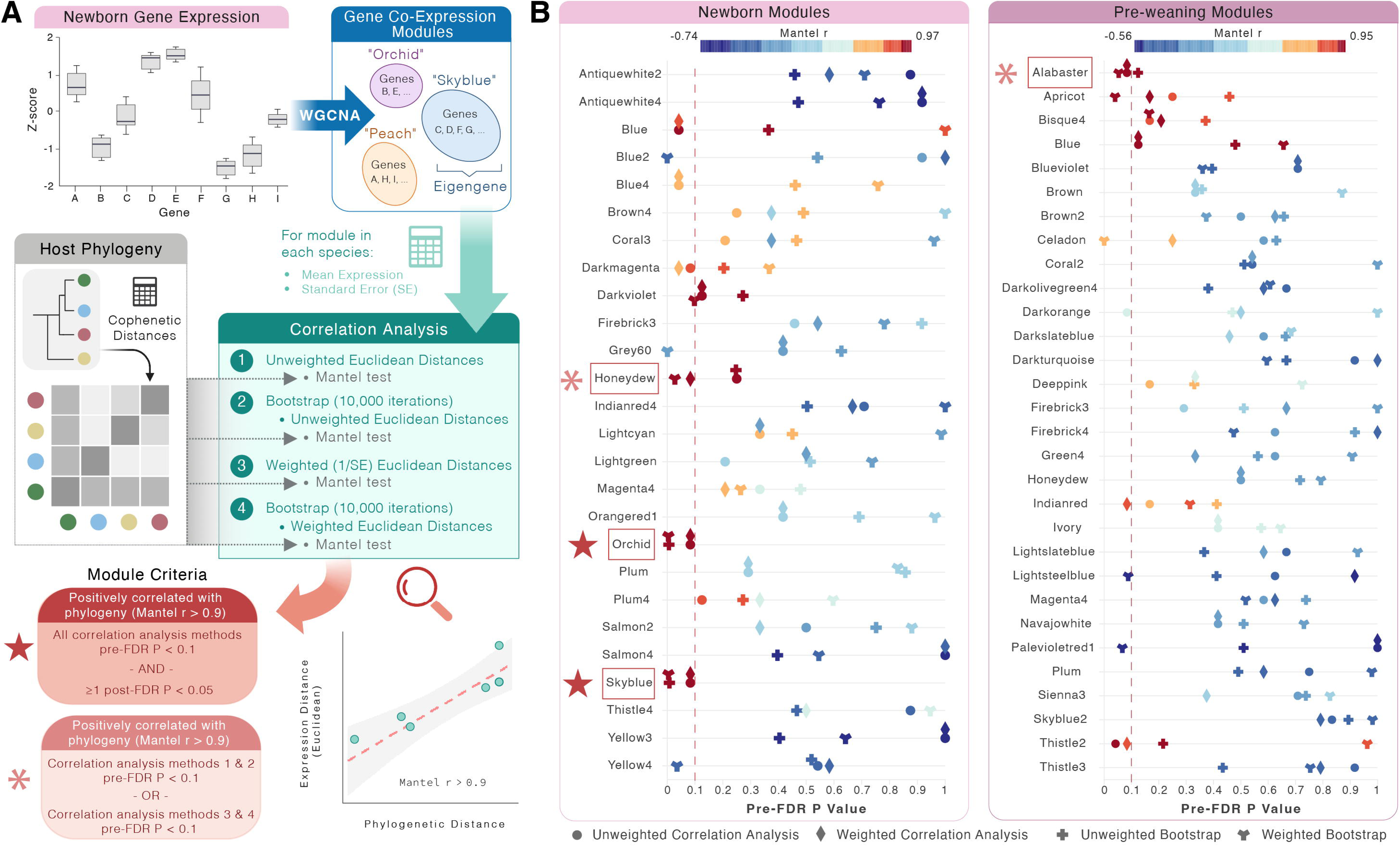
Phylogenetic correlation of weighted gene co-expression modules. **(A)** Weighted gene co-expression network analyses were conducted on expression data for each timepoint. For each module in each host species, mean expression and standard error were calculated. Correlation analyses were conducted on the host cophenetic distances and module euclidean expression distances using four methods, and modules of interest were identified based on specific criteria. **(B)** Correlation analysis results for each timepoint-specific module. Analysis method is represented by shape, while Mantel test statistic (r) is binned by color. Pre-FDR corrected P value of 0.1 is emphasized via a vertical red dashed line. Modules that match the criteria presented in **(A)** are highlighted with a red box and either an asterisk or a star. Figure 8A Created in BioRender. Rudzki, E. (2026) https://BioRender.com/ozfql43

**Fig 9.**
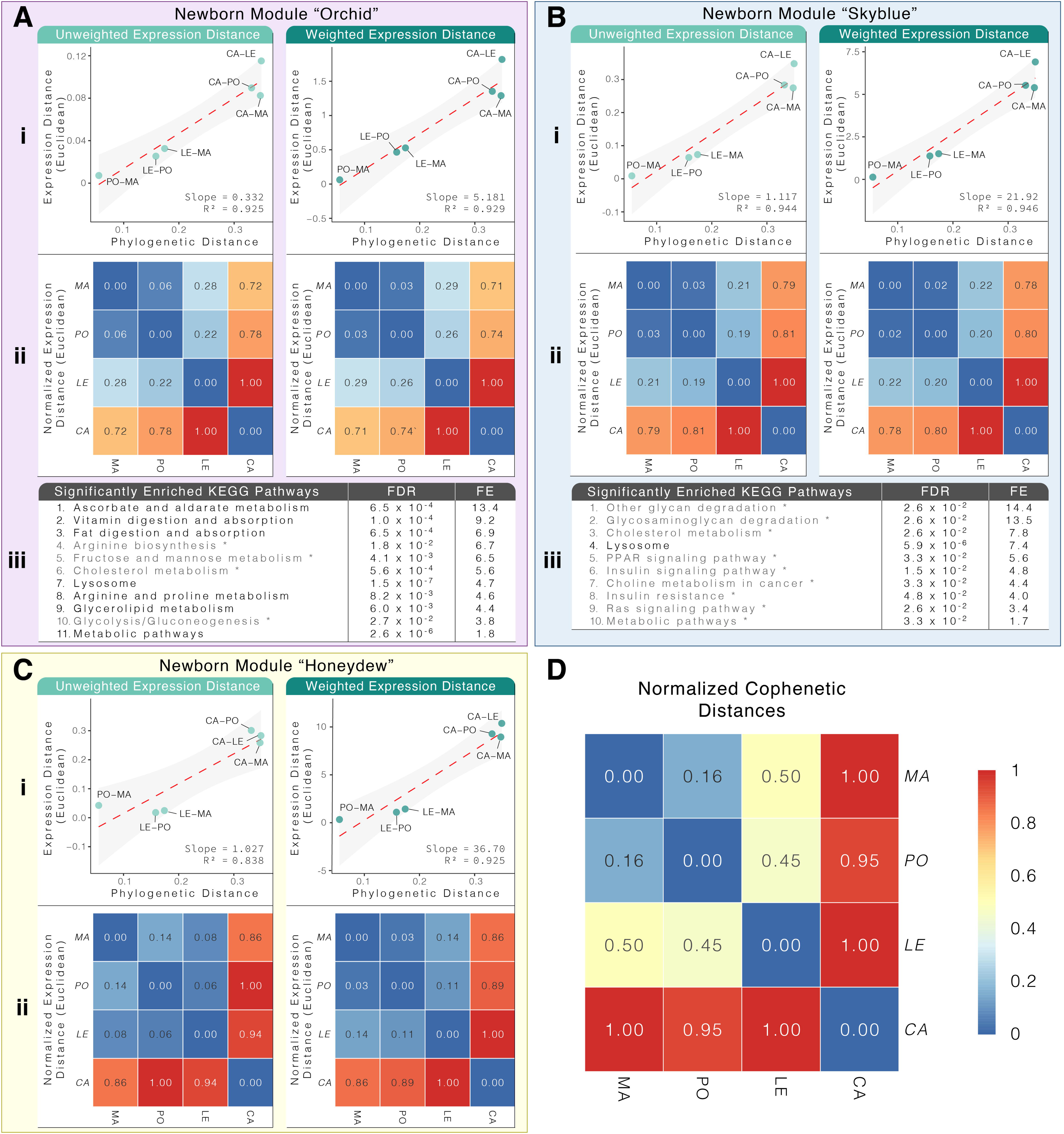
Metabolically-enriched module expression correlates strongly with *Peromyscus* phylogeny for newborn mice. Newborn timepoint modules (**A-C**) that matched the correlation criteria. **(i)** Linear regression plots of Euclidean expression distance and cophenetic distance for both unweighted (left) and weighted (right) euclidean expression distances. The slope represents the expression divergence rate of that module. Each point represents a comparison of two *Peromyscus* species, as labeled. Confidence intervals are drawn in a light grey. **(ii)** Heatmaps for unweighted (left) and weighted (right) normalized (min-max) Euclidean expression distances. **(iii)** KEGG pathways significantly enriched (FDR < 0.05) via consensus of ≥ **2** sources (functional enrichment analysis platforms ShinyGO 0.85.1 and DAVID v2025_1, and protein-protein interaction database STRING-db 12.0). Pathways noted with an asterisk (*) in grey font are conservatively considered candidate enriched pathways due to a lack of consensus between the two functional enrichment analysis platforms (FDR > 0.05 for DAVID). **(D)** Heatmap of *Peromyscus* normalized (min-max) cophenetic distances. All heatmaps in this figure are colored according to the scale shown in **D**.

### Gene modules correlated with host phylogeny at birth are enriched for metabolic pathways

Next, we performed KEGG pathway analyses to determine what pathways were enriched in each phylogenetically-correlated module. For the Orchid module, all 11 significantly enriched KEGG pathways were related to host metabolism (Fig 9Aiii). The pathways with the highest fold enrichment were ascorbate and aldarate metabolism (FE = 13.4, adj. P = 0.00065), vitamin digestion and absorption (FE = 9.2, adj. P = 0.00010), fat digestion and absorption (FE = 6.9, adj. P = 0.00065), and arginine biosynthesis* (FE = 6.7, adj. P = 0.018, pathways marked with an asterisk* are conservatively reported as candidate findings: significant in both ShinyGO and STRING-db but not confirmed across all three platforms). Notably, lysosome function was the most significantly enriched pathway in both the Orchid module (FE = 4.7, adj. P = 5.9 × 10⁻⁶) and the Skyblue module (Fig 9Biii, FE = 7.4, adj. P = 5.9 × 10⁻⁶), suggesting that lysosomal gene expression programs may be a recurring target of phylogenetic signal across independent co-expression modules. The leading pathways by fold enrichment in Skyblue included other glycan degradation* (FE = 14.4, adj. P = 0.026), glycosaminoglycan degradation* (FE = 13.5, adj. P = 0.026), and cholesterol metabolism* (FE = 7.8, adj. P = 0.026). The Honeydew module showed significant enrichment only via STRING-db ("metabolic pathways", adj. P = 0.020; "mucin type O-glycan biosynthesis", adj. P = 0.030) but not via ShinyGO or DAVID (Fig 9Ciii). The pre-weaning Alabaster module had no significantly enriched pathways. Full pathway results for all modules are provided in the S2 Appendix. The phylogenetically correlated gene modules from newborn pups were enriched for metabolic pathways, suggesting that early-life metabolic gene expression programs may be under evolutionary selection and constraint across *Peromyscus* species.

### Phylogenetically correlated gene modules show distinct rates of expression evolution

To further quantify evolutionary changes in gene module expression, we computed the slope of a linear regression between module expression and phylogenetic distance. A steeper slope reflects greater divergence in gene expression; a shallower slope indicates greater evolutionary constraint. Among the three newborn modules that met the correlation threshold, Orchid showed the slowest expression divergence rate (Fig 9Ai): Orchid diverged significantly more slowly than Skyblue (Fig 9Bi; two-tailed t-test, B-H adj. P = 0.0111 unweighted, adj. P < 0.0069 weighted) and Honeydew (Fig 9Ci; adj. P = 0.0119 unweighted, adj. P < 0.0001 weighted). Skyblue and Honeydew did not differ significantly in unweighted divergence, but Skyblue diverged more slowly than Honeydew for weighted expression (adj. P = 0.0097; Table OO in S1 Appendix). Heatmaps of normalized host cophenetic distances (Fig 9D) and Euclidean expression distances for each module (Figs 9Aii-Cii; and S5ii) illustrate how module expression divergence compares to host phylogenetic divergence.

## Discussion

A central challenge in host-microbiome dynamics is understanding how host evolutionary history influences the gut microbiota and identifying when and how these relationships are established [1, 2]. Here, using four *Peromyscus* species sampled at three developmental stages from birth through weaning, we show that phylosymbiosis in the gut microbiome and evolutionary congruence in the host transcriptome emerge at distinct, non-overlapping developmental windows. The host transcriptome is congruent with host phylogeny at birth and not thereafter, while phylosymbiosis in the gut microbiome is absent at birth and emerges only at the pre-weaning stage and is maintained thereafter. These findings reveal that evolutionary signatures in host-microbiome systems are not established simultaneously but arise through dynamic and temporally offset developmental processes and reframes phylosymbiosis as an emergent property of early mammalian development rather than a fixed feature of the adult host-microbiome system.

Several species of *Peromyscus* mice exhibited increases in diversity and substantial shifts in community structure from birth through weaning. Some elements of our data are consistent with patterns in which early composition is governed largely by vertical and contact-based maternal transmission, while later composition is increasingly shaped by host-specific selective environments. For example, the newborn microbiota was enriched for other taxa shared with the maternal mammary skin (including *Streptococcus* and a Pasteurellaceae genus) which declined as pups aged and were replaced by taxa typical of adult rodent guts, such as Muribaculaceae and Lachnospiraceae. Further, the genus *Rothia* is an oral and respiratory commensal not typically associated with the healthy mammalian gut [34, 35]. We detected *Rothia* on the mammary skin of three of our four *Peromyscus* species, and at >5% mean relative abundance in the newborn large intestine of two of those species, and it had disappeared entirely in the gut of pups by the pre-weaning timepoint. Notably, *P. polionotus* showed no *Rothia* at any body site or timepoint, showing elements of species specificity.

The newborn microbiome did not exhibit species-specific microbiomes that mirror host phylogeny (i.e., phylosymbiosis), but these patterns were observed at the pre-weaning timepoint. The delay of phylosymbiosis until the pre-weaning stage likely reflects the time required for host species-specific selective environments (shaped by traits such as gut pH, mucin composition, and immune maturation) to exert a detectable structuring influence on the colonizing community [11, 13, 15, 17, 27].

The detection of phylosymbiosis at the weaning timepoint was sensitive to the microbial dendrogram method used. The median-ceiling approach, which amplifies zero-inflation by only retaining microbiota present in at least half of the samples of a given species, yielded a significant phylosymbiosis results, while the mean-ceiling approach, which retains signal from variably present microbes, did not. This result suggests that the weaning phylosymbiotic signature is driven primarily by consistently present microbial taxa and is obscured when variable community members are included. This distinction may itself be biologically informative and could be investigated further to understand how stable or transient microbial members contribute to patterns of phylosymbiosis in the lab and in the wild [2, 36]. Most, if not all, previous phylosymbiosis studies that investigate topological congruence use a singular method of microbial table collapse [8, 37-39]. We assert that the two methods address distinctly different but related questions in the context of phylosymbiosis, and while median-ceiling fits the traditional distribution (right-skewed) of most microbiome datasets best, valuable information can be obtained by comparing both.

The congruence of host gut gene expression profiles with host phylogeny in newborns, but not at later stages, suggests a developmental window during which the intestinal transcriptome is most strongly governed by host genetic background and relatively insulated from environmental inputs, including the microbiome. Work in germ-free rodents has demonstrated that the microbiome drives broad changes in gut gene expression across immunity, metabolism, and barrier function [28, 30]. We propose that priority effects [40] are the dominant force structuring the newborn gut microbiota: the earliest colonizers arrive largely through maternal transmission, and the species-specific gut environment they encounter (shaped by divergent gene expression programs) begins to filter community composition over subsequent weeks. Under this model, birth-stage transcriptomic divergence does not directly produce phylosymbiosis, but may prime the gut to transition toward it. Consistent with this interpretation, the gene co-expression modules most strongly correlated with host phylogenetic distance at birth were enriched for metabolic pathways (e.g. lysosome function, fat and vitamin digestion and absorption, and glycan degradation) suggesting that species-specific differences in gut metabolic physiology at birth create divergent substrate and nutrient environments that differentially select for microbial taxa as development proceeds [1, 10]. The Orchid module, which exhibited the slowest rate of expression divergence among the correlated modules, may represent a core set of ancestrally constrained gut metabolic functions. As microbial colonization intensifies through the pre-weaning period, host gut gene expression is increasingly shaped by host-microbiome interactions [28, 29], which may dilute the species-specific genetic signal and erode the phylogenetic structure of the transcriptome.

Interestingly, our transcriptomic data do not strongly support the hypothesis that host immune filtering is the primary driver of phylosymbiosis. Despite the timing of phylosymbiosis emergence coinciding with the immunological weaning reaction [17, 27], the only pre-weaning gene module meeting our phylogenetic correlation criteria (Alabaster) showed no significant pathway enrichment, and dominant pre-weaning pathways across modules related to cell differentiation, proliferation, and signaling rather than immunity specifically. While the small size of the Alabaster module (78 genes, approximately half annotated) limits interpretive power, these results suggest that non-immune host filters, such as metabolic and physiological selective pressures, may play a more important role in shaping phylosymbiosis than immune-mediated selection alone [1, 3, 10].

Instead, we observe metabolic genes which underlie differences across species. For example, the expression profiles of *P. californicus* gene expression diverged most strongly, especially before weaning. This result could be due to differences in developmental trajectories: *P. californicus* pups are born markedly more precocial than the altricial young of the other three species [41] (Section A in S1 Appendix). Two gene clusters were notably elevated in *P. californicus* intestines: (1.) One cluster elevated across all timepoints contained *Slc17a3* (an organic anion transporter involved in the transport of bacterial metabolites [42]) and *Gal3st2* (a glycan sulfotransferase that catalyzes the sulfation of mucin glycans [43]). Both genes are relevant to microbial utilization of sulfated glycans as a nutrient source [44], and epithelial glycans have also been found to help induce host immune regulation of the gut microbiota [45]. Differences in the glycan composition of the intestinal mucosa may help explain why the *P. californicus* microbiota differs markedly from that of other species. (2.) A second cluster elevated in *P. californicus* specifically at weaning contained *Xab2* [46], *Top1mt* [47, 48], and *Tpt1* (orthologous to human TCTP [49]), sharing roles in nucleic acid metabolism, mitochondrial function, and cellular fitness and proliferation. The weaning-specific upregulation of this second cluster may reflect the more advanced state of intestinal maturation in *P. californicus* at this stage, given the extensive gut remodeling [50, 51] and increased metabolic demand associated with the transition from milk to solid food [52, 53].

Our study is, to our knowledge, the first to document the emergence of phylosymbiosis in early mammalian development. Further, we demonstrate that these patterns of phylosymbiosis in the gut microbiome and evolutionary congruence in the host intestinal transcriptome emerge at distinct, discordant developmental windows, indicating that host-microbiome evolutionary congruence is a dynamic rather than static property of mammalian biology. We propose that early-life host transcriptomic divergence, reflecting species-specific metabolic and physiological programs in the newborn gut, creates distinct selective environments that interact with the colonizing microbial community. Over time, these host-imposed filters drive microbial community composition toward the phylosymbiotic patterns we observe at the pre-weaning stage. As the microbiome matures and begins to reciprocally remodel host gene expression, the transcriptomic signal of host evolutionary history is progressively obscured. This model also suggests that the direction of host-microbiome influence at the gut interface may be asymmetric and dynamic across development: host-to-microbiome early in life, and microbiome-to-host during and after weaning. There may be various host-microbe interaction mechanisms that contribute to this, such as host-microbe crosstalk with bile acids [54] or microbial metabolism of host-specific milk oligosaccharides [55, 56].

These results also emphasize that early life is a critical and underexplored window for understanding the origins of host-microbiome co-evolutionary relationships [57]. Future studies could also incorporate true longitudinal designs that track the development of the microbiome in the same individuals over time. Understanding the directionality and mechanistic interactions will require experiments in which animals are inoculated with distinct microbial communities at defined developmental windows (e.g., using germ-free animals). The *Peromyscus* system, with its accessible captive stocks and growing genomic resources, is well-positioned to serve as a model for mechanistic follow-up experiments [32]. Such experiments and results will expand our understanding of how dynamic processes of microbial community assembly and mammalian gene expression underlie large-scale evolutionary patterns of microbial phylosymbiosis across host species [10].

## Materials and Methods

### 1 Animal husbandry and ethics

All mice were bred and housed in climate-controlled rooms at the Peromyscus Genetic Stock Center (PGSC) at the University of South Carolina. Animal work was approved under University of South Carolina IACUC protocol no. 2349-101211-041917. Four species were used: *P. californicus insignis* (*CA*), *P. leucopus* (*LE*), *P. polionotus subgriseus* (*PO*), and *P. maniculatus bairdii* (*MA*). Enclosures were checked once daily for new litters; more frequent checks increased stress behaviors and infanticide, especially in *PO*. Because *CA* and *PO* are monogamous and exhibit biparental care, breeding pairs of all species remained housed together throughout pup-rearing. Enclosures were cleaned twice weekly under a 16:8 light:dark cycle. Mice received *ad libitum* access to irradiated rodent chow (LabDiet Cat. No. 5L0D). Pups were weaned at species-specific timepoints (Table A in S1 Appendix).

### 2 Sample collection

Pups were sampled at three developmental timepoints: 1. birth (newborn), 2. midway through the species-specific pre-weaning period, and 3. weaning. Developmental timing and mean ages at sampling are detailed in Table A of the S1 Appendix. No more than one pup per litter was sampled at any given timepoint. At the newborn timepoint, the dam’s mammary skin was also swabbed with a sterile cotton swab (Cardinal Health Cat. No. A5000-2) for 10 seconds, then stored at −80°C. After the first pup was collected from a breeding pair, parental fecal pellets were collected at the next scheduled cleaning once the parents had been moved to a clean enclosure.

Pups were euthanized by PGSC staff using isoflurane gas followed by decapitation. Carcasses were placed in sterile sample bags (Whirlpak Cat. No. B01062), kept on ice, and transferred within 5–7 minutes. Collections were batched (1–2 newborn or pre-weaning pups, or 1–3 weaning pups at a time) to minimize time on ice, with each batch processed before the next was collected. Upon receipt, each pup was weighed and aseptically dissected (Section B1 of the S1 Appendix). The large intestine was removed, cut open laterally, placed in 500 μL sterile PBS (HyClone Cat. No. SH30258.01), and vortexed at high speed for 5 seconds. The tissue was then transferred to a second 500 μL PBS tube and vortexed again. The two PBS washes were pooled and frozen at −20°C as the “intestinal wash” fraction. The tissue was transferred to 1 mL RNAlater (Sigma Cat. No. R0901) and held at 4°C for 24 hours, after which the RNAlater was removed and the tissue stored at −80°C. All samples were shipped to the University of Pittsburgh on dry ice and stored at −80°C.

### 3 Sample processing and sequencing

#### 3.1 Microbial DNA extraction

DNA was extracted from all microbial samples (parental feces, mammary swabs, and pup intestinal washes) using the QIAamp PowerFecal DNA Isolation Kit (Qiagen Cat. No. 12830-50) per the manufacturer’s protocol. For swab samples, heads were cut off with sterile scissors and placed directly into the bead tubes. For fecal samples, portions from three pellets per parental pair were pooled for extraction. Blank extractions were included to assess kit-associated contamination [58]. DNA was stored at −80°C until sequencing.

#### 3.2 16S rRNA amplicon sequencing

The V4 region of the 16S rRNA gene was amplified using primers 515F and 806R, and resulting amplicons were sequenced on the Illumina MiSeq platform [59]. Full library preparation details are in Section B2 of the S1 Appendix. Sequence reads will be deposited in NCBI SRA upon publication.

#### 3.3 RNA extraction from large intestine tissue

Tissue was thawed on ice, finely chopped with bead-sterilized scissors, and thoroughly mixed to ensure the RNA extract represented the full length of the intestinal sample rather than any single segment. Approximately 20 mg was weighed into a sterile tube for extraction; the remainder was stored at −80°C. RNA was extracted using the RNeasy Plus Mini Kit (Qiagen Cat. No. 74134) per the manufacturer’s animal tissue protocol. Tissue was lysed in 350 μL Buffer RLT Plus (supplemented with β-mercaptoethanol at 10 μL/mL; BIO-RAD Cat. No. 1610710) by sequential trituration with a filtered P1000 tip then a sterile 22G needle/syringe (BD Cat. No. 309574), followed by centrifugation at maximum speed for 3 minutes at 4°C before proceeding to step 2 of the kit protocol. RNA quality was assessed on a 2200 TapeStation (Agilent Cat. No. G2964AA) using High Sensitivity RNA ScreenTape reagents; samples with RINe < 5 were excluded [60]. RNA was stored at −80°C until library preparation.

#### 3.4 RNA Sequencing

Libraries were prepared using the Stranded mRNA Prep, Ligation Kit (Illumina Cat. No. 20040534). The pooled library was sequenced as paired-end reads at the Children’s Hospital of Pittsburgh sequencing core. Full library preparation details are in Section B3 in the S1 Appendix. Reads will be deposited in NCBI SRA upon publication.

### 4 Data Processing and Statistical Analyses

#### 4.1 Microbiome

Raw reads were processed in QIIME2 (v2025.7) [61] using the Paired-End FASTQ Manifest method for demultiplexing and DADA2 (default settings; 220 bp trim) for denoising and amplicon sequence variant (ASV) assignment [62, 63]. A phylogenetic tree was built in QIIME2 with FastTree [64]. Taxonomy was assigned against the SILVA 138-99 reference database [65]; mitochondrial, archaeal, and chloroplast sequences were removed. Contaminant ASVs were identified and removed using the decontam package v1.22.0 [66] with the prevalence method at a threshold of 0.5 (Section B4 of the S1 Appendix). Data subsets, rarefaction depths, and excluded samples are described in Section B5 of the S1 Appendix.

Four alpha diversity metrics were calculated in QIIME2: ASV richness [62], Shannon diversity [67], Faith’s phylogenetic diversity [68], and Pielou’s evenness [69]. Kruskal-Wallis tests and Dunn’s multiple comparisons were performed in GraphPad Prism v10.6.1. Taxonomic bar plots were generated from genus-level collapsed relative abundances using phyloseq v1.46.0 [70], ggplot2 v3.5.2 [71], and associated R packages in R v4.3.3. Individual taxon abundance plots were created in Prism; Kruskal-Wallis tests and Benjamini-Hochberg (B-H) correction were performed in R. Monotonic trends were tested with the Jonckheere-Terpstra test using the DescTools package v0.99.60 [72].

Beta diversity was assessed using Jaccard (membership) and Bray-Curtis (structure) distances computed in QIIME2. Principal coordinate analysis (PCoA) plots were generated in R using ggplot2. Permutational multivariate ANOVA (adonis2, vegan v2.7-1; ‘by = margin’; 10^5^ permutations) was used to test for differences in community composition (Section B6 of the S1 Appendix). Beta dispersion was evaluated with betadisper and significance assessed by PERMADISP (permutest; [73]). Pairwise sample distances were exported from R and analyzed in Prism with Kruskal-Wallis tests and Dunn’s post-hoc comparisons. Final sample sizes are in Table B of the S1 Appendix.

#### 4.2 Host transcriptomics

Only three *Peromyscus* species had complete annotated genomes available (*P. polionotus* is incomplete); all four species were therefore mapped to the most complete available reference, *P. maniculatus* v2.1.3 (NCBI RefSeq GCF_003704035.1), selected after genome quality comparison with OMArk v0.3.0 [74]. The reference was indexed with BWA v0.7.17 [75] and a BLAST database created with BLAST+ v2.13.0 [76]. Raw reads were quality-assessed with FastQC v0.11.9 and MultiQC v1.19 [77]; base quality and adapter scores were sufficient without additional trimming. Reads were aligned with BWA-MEM, mate coordinates were checked and filled with SAMtools v1.14 [78], and alignment summaries generated with Picard v2.18.12 [79]. StringTie2 v1.3.6 [80] was used for reference-guided transcript assembly and count matrix generation.

Gene count matrices were imported into R and transformed with variance-stabilizing transformation (VST) using DESeq2 v1.42.1 [81]. PCA, Euclidean distance matrices, and adonis2 analyses were performed as described above (model equations in Section B7 of the Appendix). Beta dispersion was evaluated with vegan as previously described.

Differential gene expression was analyzed with EdgeR v4.0.16 [82]. Genes were filtered with filterByExpr (grouping by host species or timepoint as appropriate), libraries normalized, and dispersions estimated by weighted likelihood with estimateDisp (see Section B8 of the S1 Appendix for method justification). A ∼0 + *group* model matrix was fit with a negative binomial generalized log-linear model (glmQLFit) and tested by contrast (glmQLFTest). P-values were B-H adjusted; genes with FDR < 0.01 and |log2(fold change)| > 0.58 were considered differentially expressed. DEG counts were plotted with ggplot2. The top 50 highest-variance DEGs were identified, annotated with GO Biological Process terms via *Mus musculus* Entrez ID mapping (goana, limma v3.58.1 [83]), and visualized as a heatmap using ComplexHeatmap v2.18.0 [84]. Final sample sizes are in Table C of the S1 Appendix.

#### 4.3 Phylosymbiosis

Microbiome and transcriptomic data were each subset by developmental timepoint (Section B5 of the S1 Appendix). To compare transcriptomic and microbiome community structures using a common framework, the gene count matrix was converted to a BIOM file and imported into QIIME2 as a FeatureTable[Frequency] object. Features were collapsed by species using either the median-ceiling or mean-ceiling method (the former fits the typical distribution of microbiome data while the latter avoids amplifying zero-inflation; both are reported for comparison with Osborne et al. 2024 [39] and Trevelline & Moeller 2022 [38]). Bray-Curtis dendrograms were generated using UPGMA clustering [85] via qiime diversity beta-rarefaction (1,000 iterations, Spearman correlation; sampling depths in Section B5 of the S1 Appendix). Phylosymbiosis was tested following Brooks et al. 2016 [2], using ASVs rather than OTUs and the *Peromyscus* host phylogeny based on cytochrome oxidase and AVPR1A genes, pruned to the four study species.

Dendrogram congruence with the host phylogeny was assessed using Matching Cluster (MC) scores [86], which are more sensitive than Robinson-Foulds distances for closely related species as they account for branch length. Significance was evaluated as the probability of observing equal or greater congruence under stochastic assembly, based on 100,000 randomly generated bifurcating trees [87]. The limited P-value resolution is an inherent consequence of having only four species. Dendrograms were visualized with Phylo.io [88]. Rodent artwork was created with and adapted from BioRender (https://BioRender.com/pvkioap).

#### 4.4 Gene module correlation with phylogeny

To identify gene co-expression modules whose expression correlates with host phylogeny, VST-normalized counts were filtered (genes retained if ≥10 reads in ≥4 samples) and subset by timepoint. The weaning timepoint was excluded because *CA* had insufficient sample size (n = 2). Weighted gene co-expression network analysis (WGCNA v1.73; [89, 90]) was run with networkType = “signed” and corFnc = “bicor”. The host phylogenetic tree was converted to a cophenetic distance matrix (R stats v4.3.3). Mean expression and standard error per module were calculated for each species.

Module-phylogeny correlations were assessed with four methods: (1) unweighted mean-based Euclidean distances with Mantel tests (24 permutations, the maximum for four species; vegan), (2) bootstrapped version of method 1 (10,000 iterations, sampling with replacement within each species), (3) weighted mean-based Euclidean distances 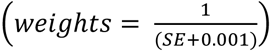 with Mantel tests, and (4) bootstrapped version of method 3. Bootstrap P-values were calculated as the proportion of resampled correlations ≥ the observed value. B-H correction was applied across all modules tested at each timepoint. The full procedure was repeated ten times across unique random seeds to confirm P-value stability.

Modules meeting either of two pre-defined criteria were considered correlated with host phylogeny: (A) Mantel r > 0.9, pre-FDR P < 0.1 for methods 1 & 2 and/or 3 & 4, and ≥1 method post-FDR P < 0.05; or (B) Mantel r > 0.9 and pre-FDR P < 0.1 for methods 1 & 2 or 3 & 4. Expression divergence rates [91] were compared across modules by ANCOVA (phylogenetic distance × module interaction), with pairwise slope differences tested by two-sided t-test and B-H correction. Cophenetic and Euclidean expression distances were min-max normalized 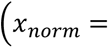 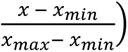 for visualization. Figures were built in R (ggplot2, pheatmap v1.0.13, ggrepel v0.9.6) and compiled in Adobe Illustrator 2026 v30.1. All custom R functions will be deposited at https://github.com/ERudzki upon publication.

#### 4.5 Module pathway enrichment analysis

Gene lists for each module were exported from the WGCNA objects; hypothetical/uncharacterized genes (LOC* identifiers) were removed. KEGG pathway enrichment was performed using three platforms: ShinyGO v0.85.1 [92], DAVID v2025_1 [93, 94], and STRING-db v12.0 [95], all with *Mus musculus* as the model organism. For ShinyGO and DAVID, module gene lists were compared against all genes across modules at that timepoint as the background. A pathway was considered significantly enriched if it met FDR < 0.05 in ≥2 of the three sources. Pathways significant in ShinyGO and STRING-db but not in DAVID are conservatively reported as candidate findings.

## Author Approvals

All authors have seen and approved the manuscript for preprint.

## Author Contributions

Elizabeth N. Rudzki – Conceptualization, data curation, formal analysis, funding acquisition, investigation, methodology, project administration, software, validation, visualization, writing – original draft preparation, writing – review & editing

Eldin Jašarević – Resources, supervision, writing – review and editing

Kevin D. Kohl – Conceptualization, funding acquisition, project administration, resources, supervision, writing – review and editing

## Abbreviations

ASV: Amplicon sequence variant
ANCOVA: Analysis of Covariance
B-H: Benjamini-Hochberg
*CA*: *Peromyscus californicus insignis*
DEGs: Differentially expressed genes
FE: Fold enrichment
*LE*: *Peromyscus leucopus*
*MA*: *Peromyscus maniculatus bairdii*
MC: Matching cluster
PCA: Principal component analysis
PCoA: Principal coordinate analysis
PGSC: Peromyscus Genetic Stock Center
*PO*: *Peromyscus polionotus subgriseus*
VST: Variance-stabilizing transformation
WGCNA: Weighted gene co-expression network analysis

## Acknowledgements

We would like to thank and acknowledge Vimala Kaza for their assistance with animal husbandry and the collection of *Peromyscus* pups, Dr. Hippokratis Kiaris for the use of their laboratory space during sample collection, and the Peromyscus Genetic Stock Center (PGSC) for temporarily housing the pups. Finally, we would like to thank José Goyco-Blas for their assistance with DNA extractions.

## Supporting Information

### Appendix. Supplemental file of appendices and tables

**S1 File. Supplemental excel file of co-expression module results.**

**Fig S1.**
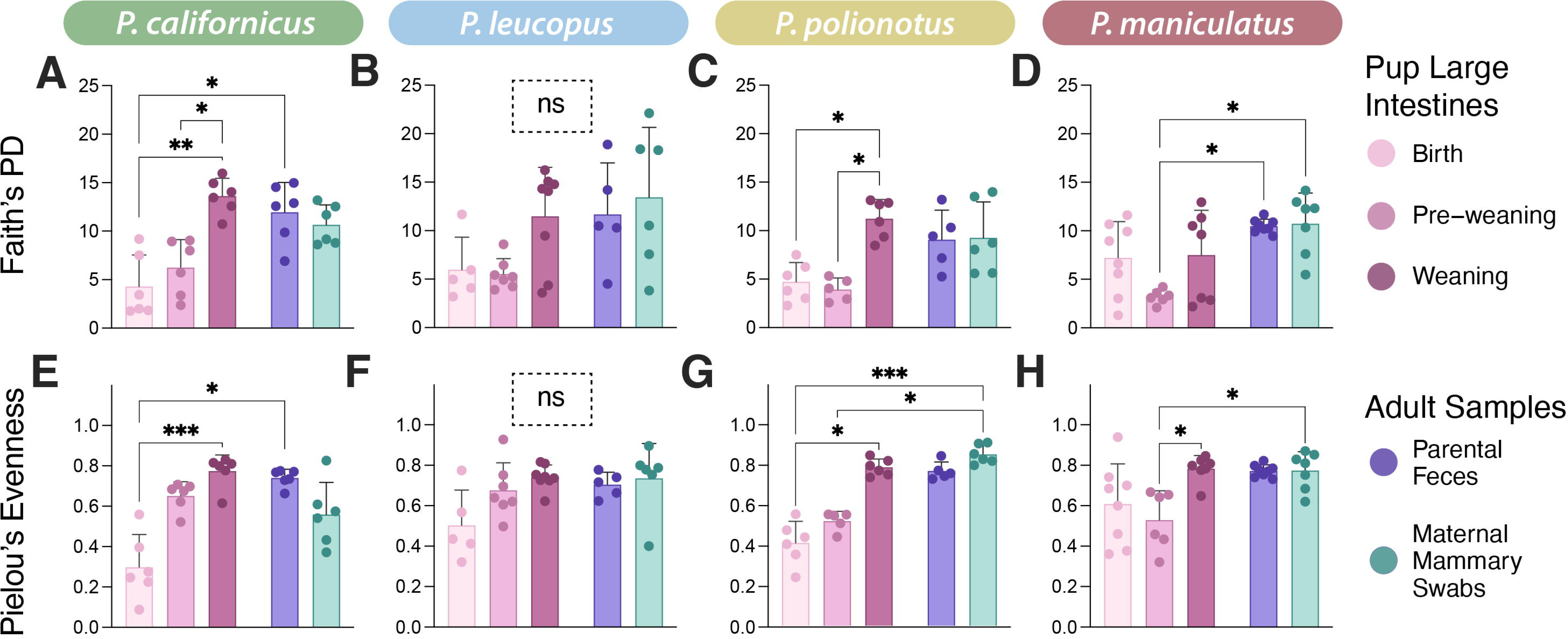
Faith’s phylogenetic diversity and Pielou’s evenness. Alpha diversity metrics of (A-D) Faith’s phylogenetic diversity and (E-H) Pielou’s evenness for each sample (represented as points). Dunn’s multiple comparison statistical significance is represented as asterisks above brackets, if present.

**Fig S2.**
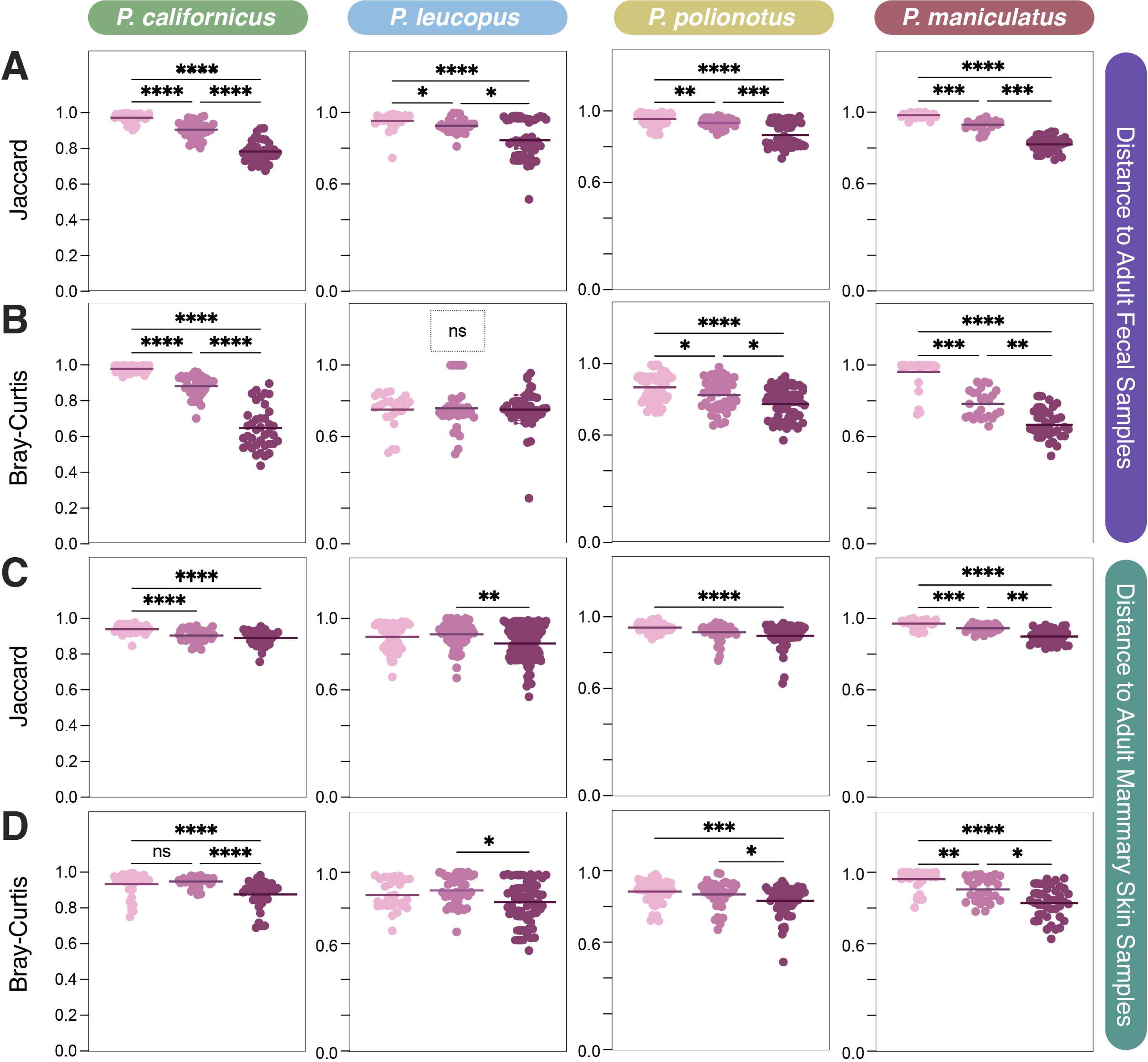
Pairwise distances comparing the microbiota of pup samples to adult samples. Pup large intestine microbiota sample distances to (A & B) adult fecal samples and (C & D) adult mammary skin samples for both Jaccard similarity (A & C) and Bray-Curtis dissimilarity (B & D). Significant Dunn’s multiple comparison test statistics are represented by asterisks above horizontal lines. Detailed statistics can be found in Tables P – U of S1 Appendix.

**Fig S3.**
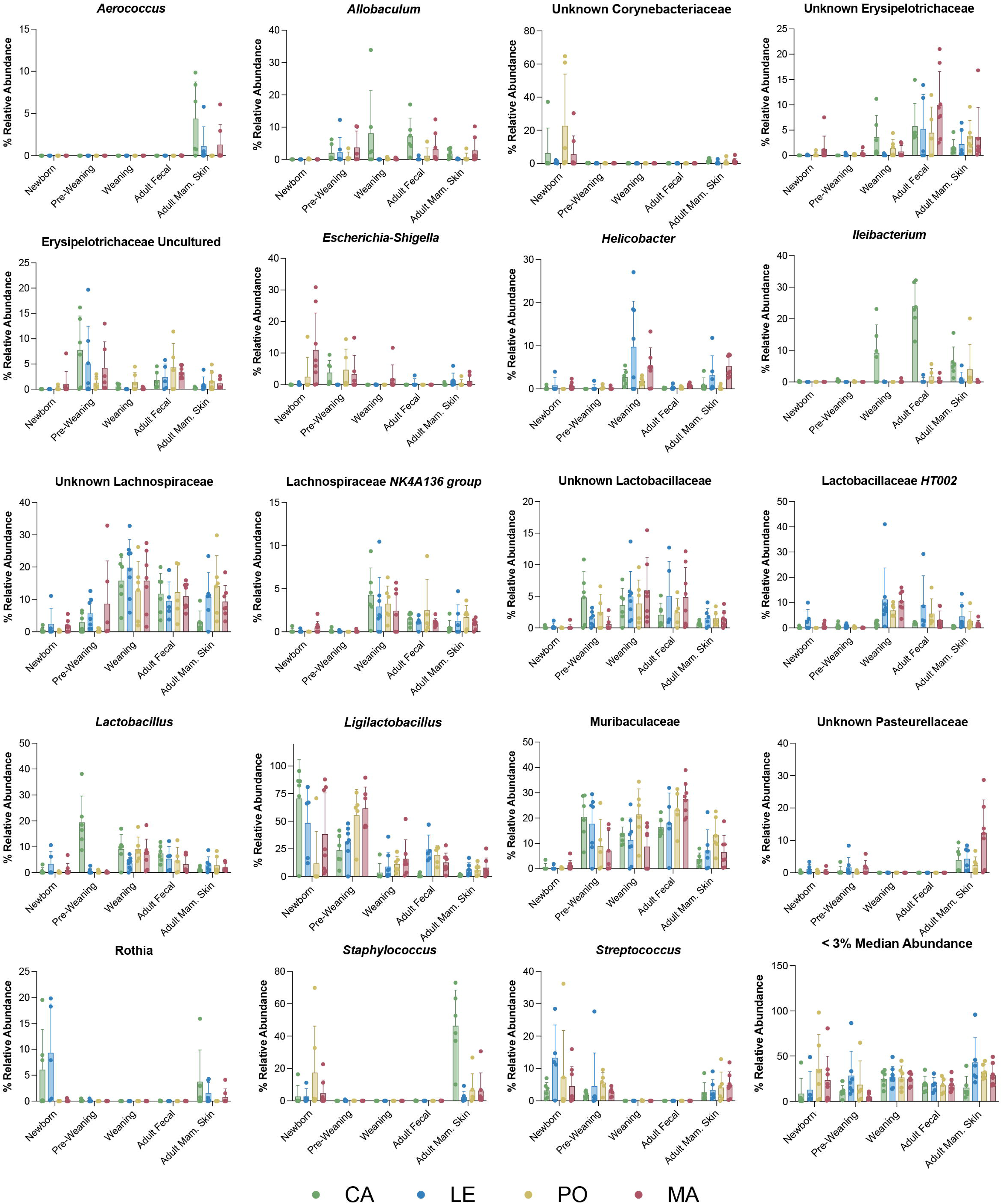
Barplots of the most abundant bacterial genera. Relative abundances of specific bacterial genera for pup intestinal wash samples (newborn, pre-weaning, and weaning timepoints), adult fecal samples, and adult mammary skin swab samples.

**Fig S4.**
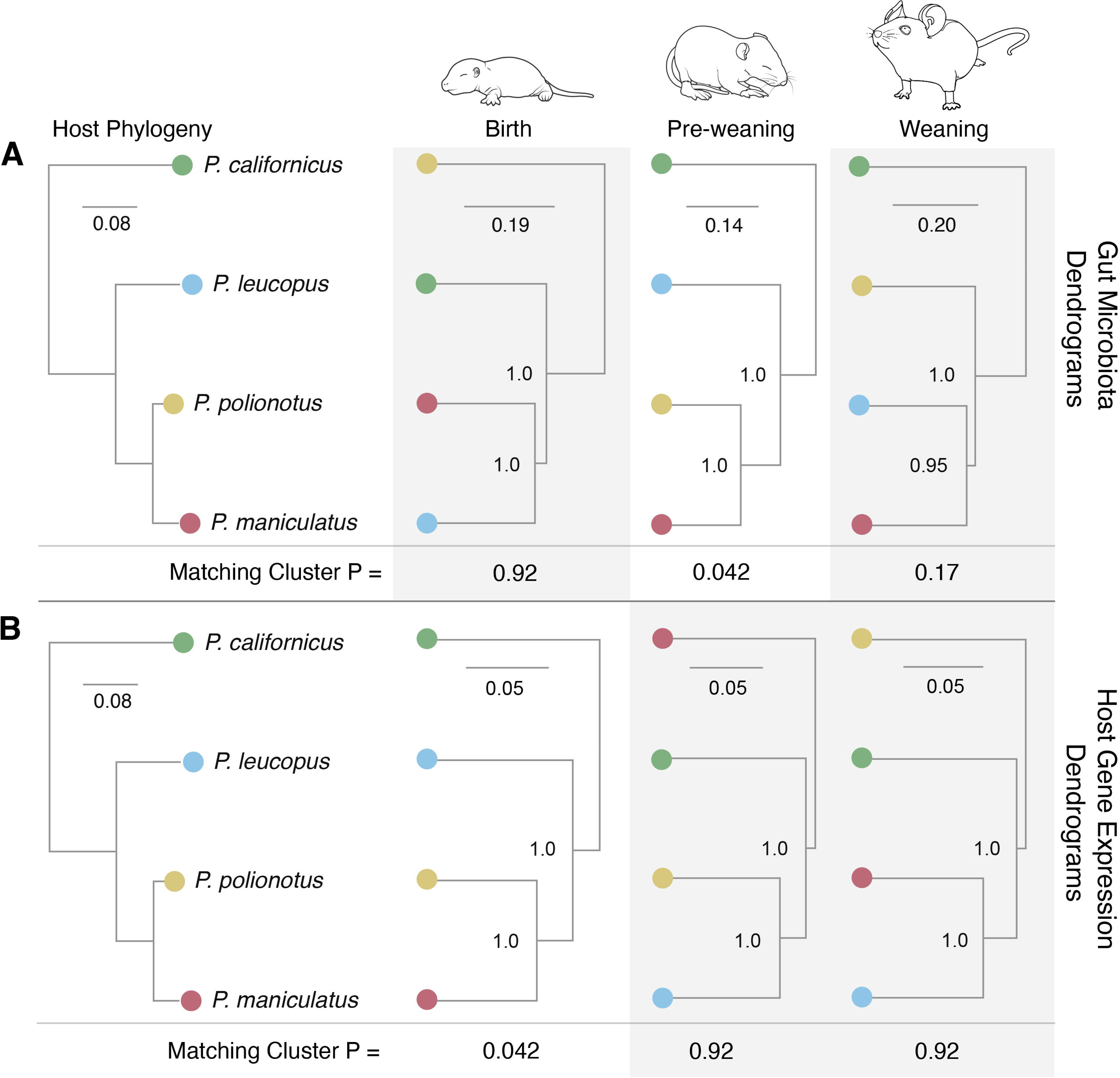
Topological congruence of Peromyscus microbiota and gene expression profiles at different stages of host development. (A) Microbiota dendrograms and (B) host gene expression dendrograms were created from Bray-Curtis dissimilarity matrices (collapsed by host species by <u>mean-ceiling</u>) and congruence compared to the host phylogenetic tree. Significance of the experimental dendrogram yielding an equal or better congruence to host phylogeny than that of 100,000 simulated random trees was calculated using matching cluster analysis. Topologically congruent dendrograms with a significant normalized matching cluster score are highlighted with a white background. Rodent artwork was created and adapted from BioRender. Rudzki, E. (2026) https://BioRender.com/pvkioap

**Fig S5.**
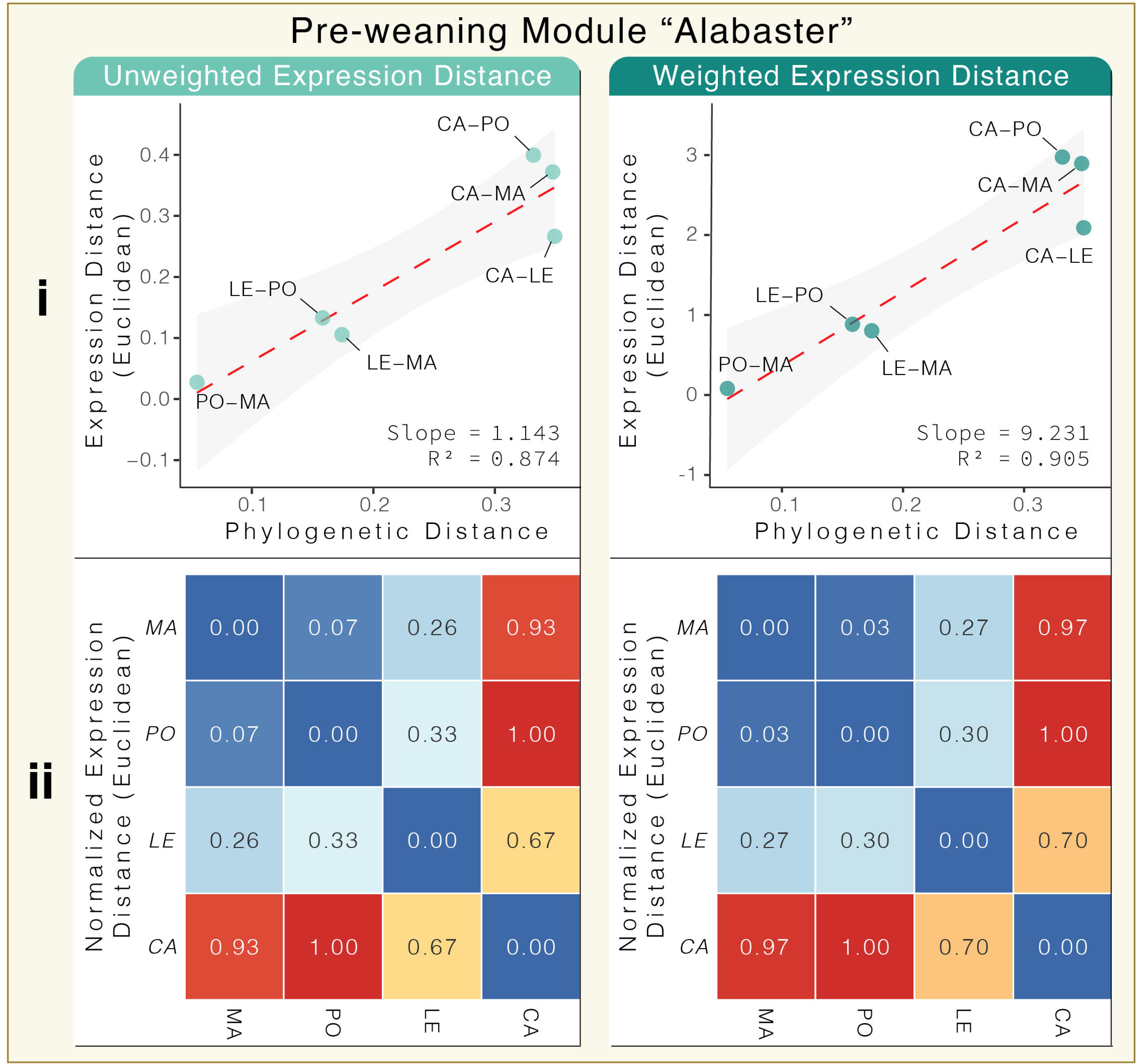
Pre-weaning timepoint module “Alabaster” that matched the correlation criteria. (i) Linear regression plots of Euclidean expression distance and cophenetic distance for both unweighted (left) and weighted (right) euclidean expression distances. The slope represents the expression divergence rate of that module. Each point represents a comparison of two *Peromyscus* species, as labeled. Confidence intervals are drawn in a light grey. (ii) Heatmaps for unweighted (left) and weighted (right) normalized (min-max) Euclidean expression distances.

