## Supplementary material for "Phylosymbiosis in the making: Discordant developmental emergence of evolutionary signatures in the mammalian gut microbiome and host transcriptome": S1 Appendix

#### Appendix A. Background

**Table A. Species specific developmental tempo.**

| Species | Body mass at adulthood | Gestation length | Weaning time | Avg age at first estrus | Lifespan in the wild |
|---|---|---|---|---|---|
| ***P. californicus*** | Mean 57 grams (Merritt 1974) | 31-33 days (Gubernick 1988) | 35-44 days (Svihla 1932; McCabe 1950) | 77 days (McCabe 1950) 44.1 days (Gubernick 1988) | At least 18 months (Merritt 1978) |
| ***P. leucopus*** | 18-23 grams (Broussard *et al.* 2009) | 22-23 days (Svihla 1932) | 22-23 days (Schmidly & Bradley 2016) | 46.22 days (Clark 1938) | Up to 18, but typically around 12 months (Schmidly & Bradley 2016) |
| ***P. polionotus*** | 10-15 grams (Wilson & Ruff 1999) | 24 days (Whitaker Jr. 1998) | 20-25 days (Wilson 1999) | 29.64 days (Clark 1938) | Around 18 months (Wilson 1999) |
| ***P. maniculatus*** | 10-24 grams (Kurta 1995) | 23-26 days (Millar & Millar 1989) | 18-24 days (King, Deshaies & Webster 1963) | 48.72 days (Clark 1938) | Less than 12 months (Baker 1983) |

Based on previous literature summarized above, we sampled pups at birth (within 24 hours). At the pre-weaning timepoint, we sampled pups at 14.3 ± 0.8 (*CA*), 13.4 ± 1.5 (*LE*), 12.6 ± 0.9 (*PO*) and 12.2 ± 1.5 (*MA*) days (mean ± sd). For the weaning timepoint, we sampled pups at 33.5 ± 5.3 (*CA*), 25.4 ± 2.1 (*LE*), 26.8 ± 1.3 (*PO*), and 23.4 ± 0.98 (*MA*) days (mean ± sd).

### Appendix B. Methods

#### Appendix B1. Aseptic dissection.

After secondary euthanasia is performed with sterilized dissection shears, the outside of the carcass is liberally sprayed with 70% ethanol and moved to a sterile petri dish. Then a midline incision is performed on the epidermis, and forceps are used to grab the exterior fur and remove the skin (via tension) from the lower abdomen. Tools are sterilized. The peritoneum is cut and dissected away to open the majority of the abdominal cavity. Tools are sterilized. Connective tissue is cut to free the stomach and intestines. The large intestine is cut 3 mm below the connection to the cecum, and about 3-5 mm above the rectum. Then the large intestine is moved to a sterile petri dish. Tools are sterilized. The large intestine is then cut laterally up the length of the organ. The tissue is then moved to the first wash tube and wash protocol proceeds as described in the main text. All tools are sterilized, the bench surface is sterilized with 70% ethanol, and gloves are changed prior to the next dissection. Multiple sets of tools are sterilizing at any given time such that tools will sit in the 70% ethanol for at least 5 minutes and then are placed in a microbead sterilizer for 60 seconds immediately prior to use. Wash tubes were pre-filled with sterile PBS under ethanol flame using irradiation-sterilized RNase- / DNase-free microcentrifuge tubes and irradiation-sterilized RNase- / DNase-free filter micropipette tips.

Appendix B2. Microbiota library preparation and amplicon sequencing methods.

Extracted DNA was sent to the University of Illinois at Chicago's DNA Services Facility for PCR amplification, library prep, and sequencing. The V4 region of the 16S rRNA gene was amplified using the 515F (GTGCCAGCMGCCGCGGTAA) and 806R (GGACTACNVGGGTWTCTAAT) primers. The forward primers contained the common sequence linker tag 1 (CS1, ACACTGACGACATGGTTCTACA) and the reverse primers contained the common sequence linker tag 2 (CS2, TACGGTAGCAGA-GACTTGGTCT) (Moonsamy *et al.* 2013). Amplicons were generated using a two-stage targeted amplicon sequencing approach (Bybee *et al.* 2011), and the resulting amplicons sequenced on the Illumina MiSeq platform (Caporaso *et al.* 2012). The first stage was performed in 96-well plates (10 µL reactions) using MyTaq HS 2X mas-termix (Bioline, London, UK) and the following thermal cycling conditions: 95ºC for 5 minutes, 28 cycles of 95ºC for 30 seconds, 55ºC for 45 seconds and 72ºC for 30 seconds. The second stage was performed in 96-well plates (20 µL reactions), also utilizing MyTaq HS 2X mas-termix. In each well a unique primer pair with a 10-base barcode was used. These were obtained from Access Array Barcode Library for Illumina (Fluidigm, South San Francisco, CA, USA). The following thermal cycling conditions were used for this second stage: 95ºC for 5 minutes, 8 cycles of 95ºC for 30 seconds, 60ºC for 30 seconds, and 72ºC for 30 seconds. A final 7-minute elongation step was performed at 72ºC.

These amplified products were pooled in equal volume using the EpMotion 5075 liquid handling robot (Eppendorf, Hamburg, Germany), and the library was purified using an AMPure XP clean up protocol (0.6X, vol/vol; Agencourt, Beckmann-Coulter, Brea, CA, USA) to remove fragments smaller than 300 base pairs. These pooled libraries, along with 20% phiX, were loaded onto an Illumina MiSeq flow cell (2x250 base paired-end reads) and sequencing primers targeting the CS1 and CS2 linker regions (Fluidigm, South San Francisco, CA, USA) were utilized to initiate sequencing.

Appendix B3. RNA library preparation and sequencing methods.

Library preparation was conducted using the Stranded mRNA Prep, Ligation Kit (Illumina Cat. No 20040534), RNA UD Indexes Set A, Ligation Kit (Illumina Cat. No. 20091655), and AMPure XP reagent (Beckman Coulter Cat. No. A63881), following manufacturer's protocol and 13 PCR cycles during library amplification. The resulting libraries were checked using the 2200 TapeStation, D5000 ScreenTape (Agilent Cat. No. 5067-5588), and D500 Reagents, Ladder, and Sample Buffer (Agilent Cat. No. 5067-5589). Libraries with sufficient amplification were pooled and sequenced on an iSeq100 using an i1 Reagent V2 (300-cycle) cartridge (Illumina Cat. No. 20031371) to verify that the libraries were pooled correctly. The pooled library (16 μL 100 ρM pooled library + 4 μL PhiX 100 ρM) was then transferred to the Health Sciences Sequencing Core at the Rangos Research Center at the Children's Hospital of Pittsburgh for RNA sequencing on the NextSeq2000 with a P4 200 flow cell.

Appendix B4. Identifying and removing contaminant ASVs.

To identify potential contaminant ASVs we used the R program 'decontam'. First, we created a 'phyloseq' object by importing the QIIME2 feature table, tree, and taxonomy using the 'qiime2R' package. DNA kit extraction blanks were first used to identify contaminants using the ‘prevalence’ method and a threshold of ‘0.5’. Fifty-six ASVs were identified as contaminants out of 3,849 unique ASVs in the dataset. Identified ASVs were then removed from the dataset in QIIME2 using the feature-table filter-features function. One sample (*PO* weaning) was excluded due to sampling depth.

Appendix B5. Description of data subsets and samples excluded.

Data sets were subset depending on the comparison or analysis needed. The microbial dataset was subset after ‘decontam’ was run and contaminant ASVs removed. The transcriptomic dataset was subset after mapping and annotation using the *P. maniculatus* reference genome (NCBI GCF_003704035.1).

##### ***Microbiota Dataset***

1. Developmental Comparisons

   DNA kit contamination control samples were removed. Data was subset by species, into 4 separate datasets (*CA, LE, MA, PO*) that each included paternal fecal samples, maternal mammary swab samples, and three timepoints (newborn, pre-weaning, and weaning) of pup large intestinal wash samples. The sampling depth threshold was set to 2300 for *CA*, 4500 for *LE* (one newborn pup intestinal wash sample was excluded), 1450 for *PO* (one weaning pup and one newborn pup intestinal samples were excluded), and 1100 for *MA* (one maternal mammary swab sample and a newborn pup intestinal wash sample were excluded). This subset was utilized for PCoA plots and statistics.

To test whether timepoint retained significance in only the pup samples, we then also removed the parental fecal samples and maternal mammary swabs from the above as well, for the purposes of ADONIS.

2. Host Species Comparisons

DNA kit contamination control samples were removed. Adult samples (fecal and maternal mammary swabs) were removed. Samples removed due to sampling depth in the Developmental Comparison (previous section) were removed. Data were then subset by timepoint into 3 separate datasets (newborn, pre-weaning, and weaning) that each included pup large intestinal wash samples from four species of *Peromyscus* mice (*CA, LE, MA, PO*). The sampling depth thresholds for each timepoint were set to 1100, 3850, and 5150, for newborn, pre-weaning, and weaning, respectively. No further samples were excluded due to sampling depth.

3. Phylosymbiosis

For topological congruence analysis, features were grouped by host species using both the median-ceiling and mean-ceiling approaches.

**Median-Ceiling:**

The species with the lowest sampling depth at each timepoint was *PO* at 1,556 for newborns, *LE* at 9,892 for pre-weaning, and *MA* at 6,917 for weaning. Rarefaction depth was determined based on the lowest sampling depth present. The sampling depth threshold was set to 1,500.

**Mean-Ceiling:**

The species with the lowest sampling depth at each timepoint was *MA* at 14,747 for newborns, *MA* at 14,693 for pre-weaning, and *MA* at 16,730 for weaning. Rarefaction depth was determined based on the lowest sampling depth present. The sampling depth threshold was set to 14,650.

4. Full Sample Set

Experimental control samples were removed and sampling depth set to 1,100 reads. Four samples were removed to match the samples excluded in the previously described datasets. This was utilized to create taxa bar plots, as well as ADONIS to test the contributions of host species and sample type (pup large intestinal wash, adult fecal, or adult mammary skin).

##### *Host Transcriptomics Dataset*

1. Full Sample Set

This sample set included all pup large intestinal tissue samples from all species and all timepoints.

2. Developmental Comparisons

Data was subset by species, into 4 separate datasets (*CA*, *LE*, *MA*, *PO*) that each included three timepoints (newborn, pre-weaning, and weaning) of pup large intestinal tissue samples.

3. Host Species Comparisons

Data was subset by timepoint, into 3 separate datasets (newborn, pre-weaning, and weaning) that each included pup large intestinal tissue samples from four species of *Peromyscus* mice (*CA*, *LE*, *PO*, *MA*).

4. Phylosymbiosis

For topological congruence analysis, features were grouped by host species using both the median-ceiling and mean-ceiling approaches.

**Median-Ceiling:**

The species with the lowest sampling depth at each timepoint was *CA* at 37,291,666 for newborns, *CA* at 34,503,290 for pre-weaning, and *CA* at 31,913,566 for weaning. Rarefaction depth was determined based on the lowest sampling depth present. The sampling depth threshold was set to 31,900,000.

**Mean-Ceiling:**

The species with the lowest sampling depth at each timepoint was *LE* at 39,777,354 for newborns, *CA* at 36,672,425 for pre-weaning, and *CA* at 31,913,566 for weaning. Rarefaction depth was determined based on the lowest sampling depth present. The sampling depth threshold was set to 31,900,000.

Appendix B6. Example adonis and betadisper formulas utilized for the microbiota data subsets.

***Developmental Comparisons***

Full sample set:

Permutation test for adonis under reduced model:

```
adonis2(formula = jaccard ~ Species + Type, data = metadata, permutations = 1e+05,
by = "margin")
```

Wherein `Type` is a factor of 3 levels: adult fecal, adult mammary skin swab, pup large intestinal wash, and `Species` is a factor of 4 levels: *CA*, *LE*, *PO*, and *MA*.

Only pup (large intestinal wash) samples, subset by species:

Permutation test for adonis under reduced model

```
adonis2(formula = jaccard_CA_pup ~ Timepoint_Cat + CageID, data = metadata_CA_pup,
permutations = 1e+05, by = "margin")
```

Wherein `Timepoint_Cat` is a factor of 3 levels: newborn, pre-weaning, and weaning.

Homogeneity of multivariate dispersions

```
betadisper(d = jaccard_CA_pup, group = CA_fac, type = c("centroid"), bias.adjust
= FALSE, sqrt.dist = FALSE, add = FALSE)
```

Wherein `group` is a factor of 3 levels: newborn, pre-weaning, and weaning.

***Species Comparisons***

Only pup (large intestinal wash) samples, subset by timepoint:

Permutation test for adonis under reduced model

```
adonis2(formula = jaccard_TP1 ~ Species, data = metadata_TP1, permutations =
1e+05, by = "margin")
```

Wherein `Species` is a factor of 4 levels*: CA, LE, PO*, and *MA*.

Homogeneity of multivariate dispersions

```
betadisper(d = jaccard_TP1, group = TP1_fac, type = c("centroid"), bias.adjust =
FALSE, sqrt.dist = FALSE, add = FALSE)
```

Wherein the group is a factor of 4 levels: *CA, LE, PO*, and *MA*.

Appendix B7. Example adonis and betadisper formulas utilized for the transcriptomics data subsets.

***Developmental Comparisons***

Pup (large intestine tissue) samples, subset by species:

Permutation test for adonis under reduced model

```
adonis2(formula = dist_matrix_CA ~ Timepoint_Cat, data = pcaData_CA, permutations
= 1e+05, by = "margin")
```

Wherein `Timepoint_Cat` is a factor of 3 levels: newborn, pre-weaning, and weaning.

Homogeneity of multivariate dispersions

```
betadisper(d = dist_matrix_CA, group = pcaData_CA$Timepoint, type = c("centroid"),
bias.adjust = FALSE, sqrt.dist = FALSE, add = FALSE)
```

Wherein `group` is a factor of 3 levels: newborn, pre-weaning, and weaning.

***Species Comparisons***

Pup (large intestine tissue) samples, subset by timepoint.

Permutation test for adonis under reduced model

```
adonis2(formula = dist_matrix_TP1 ~ Species, data = pcaData_TP1, permutations =
1e+05, by = "margin")
```

Wherein `Species` is a factor of 4 levels: *CA, LE, PO*, and *MA*.

Homogeneity of multivariate dispersions

```
betadisper(d = dist_matrix_TP1, group = pcaData_TP1$Species, type = c("centroid"),
bias.adjust = FALSE, sqrt.dist = FALSE, add = FALSE)
```

Wherein group is a factor of 4 levels: *CA, LE, PO*, and *MA*.

Appendix B8. EdgeR dispersion comparison.

Our dataset is a longitudinal study across host development, and each pup was sampled only once and thus is represented at a singular timepoint. While we do have replicates of each species at each timepoint, we wanted to compare two different estimation of dispersion methods in EdgeR: one that is the general function used for experiments with biological replicates (‘estimateDisp’), and one that is a specialized suggestion for experiments without biological replicates (‘estimateGLMCommonDisp’ with ‘method’ = “deviance”, ‘robust’ = “TRUE”, and ‘subset’ = “NULL”). Both methods resulted in a BCV around 0.31 $\pm$ ~0.025. The typical expected values of the square root dispersion (BCV) are about 0.4 for controlled experiments on populations such as human datasets, or about 0.1 for data on genetically identical model organisms (Robinson, McCarthy & Smyth 2010). Given this, our result of ~0.3 is reasonable for a study involving four species of mice that have been bred captively to retain the genetic variation of their original sampled populations. As both methods produced similar values, we moved forward with the method that consistently produced the best visual trend fit with the data, which was ‘estimateDisp’.

#### Appendix C. Tables

**Table B. Microbiome dataset sample distribution.**

Final sample sizes for the 16S amplicon (microbiome) dataset.

| | | Feces | Mam.Swab | Large Intestine Wash | | |
|---|---|---|---|---|---|---|
| | Total | Adult | Adult | Newborn Pups | Pre-weaning Pups | Weaning Pups |
| CA | 30 | 6 | 6 | 6 | 6 | 6 |
| LE | 31 | 5 | 6 | 5 | 7 | 8 |
| PO | 28 | 5 | 6 | 6 | 5 | 6 |
| MA | 36 | 8 | 7 | 8 | 6 | 7 |

**Table C. Host transcriptomic dataset sample distribution.**

Final sample sizes for the RNAseq (transcriptomics) dataset.

| | | Large Intestine Wash | | |
|---|---|---|---|---|
| | Total | Newborn Pups | Pre-weaning Pups | Weaning Pups |
| CA | 11 | 5 | 4 | 2 |
| LE | 17 | 4 | 7 | 6 |
| PO | 13 | 6 | 4 | 3 |
| MA | 15 | 6 | 4 | 5 |

Appendix C1. Microbial alpha diversity – Developmental comparison.

**Table D. Kruskal-Wallis tests for microbial alpha diversity.**

Kruskal-Wallis statistics on the developmental comparison data subsets (*CA*, *LE*, *PO*, and *MA*) for alpha diversity metrics of #ASVs (community richness), Shannon diversity index, Faith's phylogenetic diversity, and Pielou's evenness, across 5 groups: adult fecal samples, adult mammary swab samples, and newborn, pre-weaning, and weaning pup large intestinal wash samples.

| Subset | Alpha Diversity Metric | Kruskal-Wallis P | Number of Groups | Kruskal-Wallis Statistic |
|---|---|---|---|---|
| *CA* | # ASVs | **0.0007** | 5 | 19.4 |
| | Shannon Diversity | **0.0002** | 5 | 21.8 |
| | Faith's PD | **0.0007** | 5 | 19.3 |
| | Pielou's Evenness | **0.0006** | 5 | 19.5 |
| *LE* | # ASVs | 0.125 | 5 | 7.21 |
| | Shannon Diversity | **0.0394** | 5 | 10.1 |
| | Faith's PD | 0.0821 | 5 | 8.27 |
| | Pielou's Evenness | 0.0733 | 5 | 8.56 |
| *PO* | # ASVs | **0.0006** | 5 | 19.5 |
| | Shannon Diversity | **0.0005** | 5 | 20.0 |
| | Faith's PD | **0.0021** | 5 | 16.8 |
| | Pielou's Evenness | **0.0002** | 5 | 22.3 |
| *MA* | # ASVs | **0.0040** | 5 | 15.4 |
| | Shannon Diversity | **0.0012** | 5 | 18.0 |
| | Faith's PD | **0.0091** | 5 | 13.5 |
| | Pielou's Evenness | **0.0039** | 5 | 15.4 |

**Table E. Dunn's multiple comparisons test: #ASVs.**

Dunn's multiple comparisons tests for the number of unique ASVs (community richness) for the developmental comparison data subsets (*CA*, *LE*, *PO*, and *MA*). The groups being compared are listed in the first two columns of each row, and are adult fecal samples, adult mammary swab samples, or newborn, pre-weaning, or weaning pup large intestinal wash samples.

| | | *CA* | | *LE* | | *PO* | | *MA* | |
|---|---|---|---|---|---|---|---|---|---|
| Group 1 | Group 2 | Mean rank diff. | Adj. P Value | Mean rank diff. | Adj. P Value | Mean rank diff. | Adj. P Value | Mean rank diff. | Adj. P Value |
| Par. Feces | Mam. Swabs | 5.33 | >0.99 | 0.133 | >0.99 | 2.35 | >0.99 | 2.85 | >0.99 |
| Par. Feces | Newborn | 15.0 | **0.0317** | 10.7 | 0.627 | 13.0 | 0.0893 | 14.1 | 0.0731 |
| Par. Feces | Pre-weaning | 10.3 | 0.421 | 8.16 | >0.99 | 15.0 | **0.0392** | 18.7 | **0.0099** |
| Par. Feces | Weaning | -4.83 | >0.99 | 0.800 | >0.99 | -1.73 | >0.99 | 6.42 | >0.99 |
| Mam. Swabs | Newborn | 9.67 | 0.572 | 10.6 | 0.549 | 10.7 | 0.246 | 11.3 | 0.3857 |
| Mam. Swabs | Pre-weaning | 5.00 | >0.99 | 8.02 | >0.99 | 12.7 | 0.111 | 15.9 | 0.0672 |
| Mam. Swabs | Weaning | -10.2 | 0.455 | 0.667 | >0.99 | -4.08 | >0.99 | 3.57 | >0.99 |
| Newborn | Pre-weaning | -4.67 | >0.99 | -2.54 | >0.99 | 1.98 | >0.99 | 4.60 | >0.99 |
| Newborn | Weaning | -19.8 | **0.0010** | -9.90 | 0.560 | -14.8 | **0.0189** | -7.71 | >0.99 |
| Pre-weaning | Weaning | -15.2 | **0.0284** | -7.36 | >0.99 | -16.7 | **0.0078** | -12.3 | 0.3566 |

**Table F. Dunn's multiple comparisons test: Shannon Diversity.**

Dunn's multiple comparisons tests for Shannon diversity for the developmental comparison data subsets (*CA*, *LE*, *PO*, and *MA*). The groups being compared are listed in the first two columns of each row, and are adult fecal samples, adult mammary swab samples, or newborn, pre-weaning, or weaning pup large intestinal wash samples.

| | | *CA* | | *LE* | | *PO* | | *MA* | |
|---|---|---|---|---|---|---|---|---|---|
| Group 1 | Group 2 | Mean rank diff. | Adj. P Value | Mean rank diff. | Adj. P Value | Mean rank diff. | Adj. P Value | Mean rank diff. | Adj. P Value |
| Par. Feces | Mam. Swabs | 9.33 | 0.663 | -2.97 | >0.99 | -2.30 | >0.99 | 0.375 | >0.99 |
| Par. Feces | Newborn | 17.7 | **0.0051** | 11.4 | 0.474 | 13.4 | 0.0729 | 14.0 | 0.0787 |
| Par. Feces | Pre-weaning | 8.33 | >0.99 | 6.06 | >0.99 | 10.8 | 0.379 | 18.5 | **0.0112** |
| Par. Feces | Weaning | -3.67 | >0.99 | -1.68 | >0.99 | -2.80 | >0.99 | 3.09 | >0.99 |
| Mam. Swabs | Newborn | 8.33 | >0.99 | 14.4 | 0.0907 | 15.7 | **0.0097** | 13.6 | 0.125 |
| Mam. Swabs | Pre-weaning | -1.00 | >0.99 | 9.02 | 0.744 | 13.1 | 0.0854 | 18.2 | **0.0194** |
| Mam. Swabs | Weaning | -13.0 | 0.105 | 1.29 | >0.99 | -0.500 | >0.99 | 2.71 | >0.99 |
| Newborn | Pre-weaning | -9.33 | 0.663 | -5.34 | >0.99 | -2.57 | >0.99 | 4.54 | >0.99 |
| Newborn | Weaning | -21.3 | **0.0003** | -13.1 | 0.117 | -16.2 | **0.0066** | -10.9 | 0.454 |
| Pre-weaning | Weaning | -12.0 | 0.182 | -7.73 | >0.99 | -13.6 | 0.0633 | -15.5 | 0.0838 |

**Table G. Dunn's multiple comparisons test: Faith's Phylogenetic Diversity.**

Dunn's multiple comparisons tests for Faith's phylogenetic diversity for the developmental comparison data subsets (*CA*, *LE*, *PO*, and *MA*). The groups being compared are listed in the first two columns of each row, and are adult fecal samples, adult mammary swab samples, or newborn, pre-weaning, or weaning pup large intestinal wash samples.

| | | *CA* | | *LE* | | *PO* | | *MA* | |
|---|---|---|---|---|---|---|---|---|---|
| Group 1 | Group 2 | Mean rank diff. | Adj. P Value | Mean rank diff. | Adj. P Value | Mean rank diff. | Adj. P Value | Mean rank diff. | Adj. P Value |
| Par. Feces | Mam. Swabs | 4.00 | >0.99 | -1.40 | >0.99 | -0.867 | >0.99 | -2.82 | >0.99 |
| Par. Feces | Newborn | 14.7 | **0.0391** | 9.60 | 0.950 | 9.80 | 0.491 | 8.00 | >0.99 |
| Par. Feces | Pre-weaning | 12.0 | 0.182 | 9.31 | 0.802 | 12.4 | 0.172 | 16.4 | **0.0391** |
| Par. Feces | Weaning | -4.00 | >0.99 | 0.850 | >0.99 | -3.87 | >0.99 | 6.61 | >0.99 |
| Mam. Swabs | Newborn | 10.7 | 0.359 | 11.0 | 0.457 | 10.7 | 0.247 | 10.8 | 0.472 |
| Mam. Swabs | Pre-weaning | 8.00 | >0.99 | 10.7 | 0.342 | 13.3 | 0.0774 | 19.2 | **0.0103** |
| Mam. Swabs | Weaning | -8.00 | >0.99 | 2.25 | >0.99 | -3.00 | >0.99 | 9.43 | 0.941 |
| Newborn | Pre-weaning | -2.67 | >0.99 | -0.286 | >0.99 | 2.60 | >0.99 | 8.42 | >0.99 |
| Newborn | Weaning | -18.7 | **0.0024** | -8.75 | 0.914 | -13.7 | **0.0401** | -1.39 | >0.99 |
| Pre-weaning | Weaning | -16.0 | **0.0164** | -8.46 | 0.721 | -16.3 | **0.0109** | -9.81 | 0.942 |

**Table H. Dunn's multiple comparisons test: Pielou's Evenness.**

Dunn's multiple comparisons tests for Pielou's evenness for the developmental comparison data subsets (*CA*, *LE*, *MA*, and *PO*). The groups being compared are listed in the first two columns of each row, and are adult fecal samples, adult mammary swab samples, or newborn, pre-weaning, weaning, post-weaning pup large intestinal wash samples.

| | | *CA* | | *LE* | | *PO* | | *MA* | |
|---|---|---|---|---|---|---|---|---|---|
| Group 1 | Group 2 | Mean rank diff. | Adj. P Value | Mean rank diff. | Adj. P Value | Mean rank diff. | Adj. P Value | Mean rank diff. | Adj. P Value |
| Par. Feces | Mam. Swabs | 8.67 | 0.882 | -6.23 | >0.99 | -7.57 | >0.99 | -2.04 | >0.99 |
| Par. Feces | Newborn | 16.5 | **0.0117** | 8.20 | >0.99 | 12.1 | 0.151 | 9.63 | 0.677 |
| Par. Feces | Pre-weaning | 5.50 | >0.99 | 2.03 | >0.99 | 8.80 | 0.908 | 15.3 | 0.0736 |
| Par. Feces | Weaning | -4.00 | >0.99 | -3.78 | >0.99 | -2.07 | >0.99 | -2.75 | >0.99 |
| Mam. Swabs | Newborn | 7.83 | >0.99 | 14.4 | 0.0875 | 19.7 | **0.0003** | 11.7 | 0.325 |
| Mam. Swabs | Pre-weaning | -3.17 | >0.99 | 8.26 | >0.99 | 16.4 | **0.0102** | 17.3 | **0.0319** |
| Mam. Swabs | Weaning | -12.7 | 0.1270 | 2.46 | >0.99 | 5.50 | >0.99 | -0.714 | >0.99 |
| Newborn | Pre-weaning | -11.0 | 0.3045 | -6.17 | >0.99 | -3.30 | >0.99 | 5.63 | >0.99 |
| Newborn | Weaning | -20.5 | **0.0005** | -12.0 | 0.209 | -14.2 | **0.0286** | -12.4 | 0.232 |
| Pre-weaning | Weaning | -9.50 | 0.6161 | -5.80 | >0.99 | -10.9 | 0.291 | -18.0 | **0.0213** |

Appendix C2. Microbial beta diversity – Full sample set.

**Table I. ADONIS results for full microbiota sample set.**

ADONIS statistics on Jaccard and Bray-Curtis distances for the full sample set wherein species is *CA*, *LE*, *PO*, and *MA*, and sample type is pup large intestinal washes, adult parental feces, and adult maternal mammary swabs. Adonis2 formula of Jaccard ~ HostSpecies + SampleType, with 1 x $10^5$ permutations and "by = margin". Both sample type and host species explain a significant amount of the beta diversity in the sample set, however the type of sample has a ~1.5-2x larger effect size than host species.

| Subset | | Df | SumofSqs | $R^2$ | F | Pr(>F) |
|---|---|---|---|---|---|---|
| *Jaccard* | Species | 3 | 2.8 | 0.0541 | 2.41 | 1.0 x $10^{-5}$ |
| | SampleType | 2 | 2.8 | 0.0541 | 3.61 | 1.0 x $10^{-5}$ |
| | Residual | 119 | 47 | 0.892 | | |
| *Bray-Curtis* | Species | 3 | 3.4 | 0.0762 | 3.68 | 1.0 x $10^{-5}$ |
| | SampleType | 2 | 4.6 | 0.102 | 7.39 | 1.0 x $10^{-5}$ |
| | Residual | 119 | 37 | 0.821 | | |

Appendix C3. Microbial beta diversity – Developmental comparison.

**Table J. ADONIS results for pup samples, Jaccard similarity.**

ADONIS statistics on Jaccard distances for the developmental comparison of pup (large intestinal wash) samples. Adonis2 formula of Jaccard ~ Timepoint + CageID, with 1 x $10^5$ permutations and "by = margin". Developmental timepoint explains a significant amount of beta diversity variation in all *Peromyscus* species.

| Subset | | Df | SumofSqs | $R^2$ | F | Pr(>F) |
|---|---|---|---|---|---|---|
| *CA* | Timepoint | 1 | 0.71 | 0.102 | 1.81 | **0.0268** |
| | CageID | 10 | 3.1 | 0.451 | 0.800 | 0.969 |
| | Residual | 5 | 2.0 | 0.282 | | |
| *LE* | Timepoint | 2 | 1.2 | 0.154 | 1.67 | **0.00332** |
| | CageID | 8 | 3.2 | 0.397 | 1.07 | 0.239 |
| | Residual | 9 | 3.3 | 0.416 | | |
| *PO* | Timepoint | 2 | 1.4 | 0.211 | 1.94 | **0.00027** |
| | CageID | 7 | 2.4 | 0.362 | 0.952 | 0.668 |
| | Residual | 7 | 2.6 | 0.380 | | |
| *MA* | Timepoint | 2 | 1.1 | 0.139 | 1.58 | **0.0075** |
| | CageID | 9 | 3.6 | 0.439 | 1.11 | 0.156 |
| | Residual | 9 | 3.2 | 0.396 | | |

**Table K. Average distance to centroid for Jaccard similarity.**

Beta dispersion estimates on Jaccard distances for the developmental comparison of pup (large intestinal wash) samples. Average distance to the centroid for each species and timepoint is reported.

| | Large Intestine Wash | | |
|---|---|---|---|
| | Newborn Pups | Pre-weaning Pups | Weaning Pups |
| CA | 0.557 | 0.527 | 0.502 |
| LE | 0.560 | 0.576 | 0.560 |
| PO | 0.602 | 0.500 | 0.514 |
| MA | 0.616 | 0.506 | 0.562 |

**Table L. Permutation test for homogeniety of multivariate dispersions for Jaccard similarity.**

Beta dispersion significance for the Jaccard distances of the developmental comparison of pup (large intestinal wash) samples. *P. polionotus* and *P. maniculatus* samples differ in dispersion by timepoint for community membership.

| Subset | | Df | SumofSqs | Mean Sq | F value | N.Perm | Pr(>F) |
|---|---|---|---|---|---|---|---|
| CA | Timepoint | 2 | 0.0093 | 0.0046 | 1.26 | 99 | 0.39 |
| | Residuals | 15 | 0.055 | 0.0037 | | | |
| LE | Timepoint | 2 | 0.0012 | 0.00059 | 0.120 | 99 | 0.94 |
| | Residuals | 17 | 0.083 | 0.0049 | | | |
| PO | Timepoint | 2 | 0.035 | 0.018 | 27.8 | 99 | **0.01** |
| | Residuals | 14 | 0.0088 | 0.00063 | | | |
| MA | Timepoint | 2 | 0.042 | 0.021 | 14.7 | 99 | **0.01** |
| | Residuals | 18 | 0.026 | 0.0014 | | | |

**Table M. ADONIS results for pup samples, Bray-Curtis dissimilarity.**

ADONIS statistics on Bray-Curtis distances for the developmental comparison of pup (large intestinal wash) samples. Adonis2 formula of braycurtis ~ Timepoint + CageID, with 1 x $10^5$ permutations and "by = margin". Developmental timepoint explains a significant amount of beta diversity variation in all *Peromyscus* species.

| Subset | | Df | SumofSqs | $R^2$ | F | Pr(>F) |
|---|---|---|---|---|---|---|
| *CA* | Timepoint | 1 | 0.94 | 0.165 | 4.30 | **0.0045** |
| | CageID | 10 | 2.1 | 0.368 | 0.960 | 0.56 |
| | Residual | 5 | 1.1 | 0.192 | | |
| *LE* | Timepoint | 2 | 1.4 | 0.225 | 2.94 | **0.00061** |
| | CageID | 8 | 2.3 | 0.364 | 1.19 | 0.193 |
| | Residual | 9 | 2.2 | 0.343 | | |
| *PO* | Timepoint | 2 | 1.7 | 0.289 | 3.25 | **0.00131** |
| | CageID | 7 | 1.8 | 0.304 | 0.978 | 0.519 |
| | Residual | 7 | 1.8 | 0.311 | | |
| *MA* | Timepoint | 2 | 1.4 | 0.240 | 3.24 | **0.00099** |
| | CageID | 9 | 2.2 | 0.372 | 1.12 | 0.311 |
| | Residual | 9 | 2.0 | 0.333 | | |

**Table N. Average distance to centroid for Bray-Curtis dissimilarity.**

Beta dispersion estimates on Bray-Curtis distances for the developmental comparison of pup (large intestinal wash) samples. Average distance to the centroid for each species and timepoint is reported.

| | Large Intestine Wash | | |
|---|---|---|---|
| | Newborn Pups | Pre-weaning Pups | Weaning Pups |
| CA | 0.300 | 0.417 | 0.451 |
| LE | 0.367 | 0.457 | 0.504 |
| PO | 0.575 | 0.310 | 0.414 |
| MA | 0.516 | 0.294 | 0.463 |

**Table O. Permutation test for homogeneity of multivariate dispersions for Bray-Curtis dissimilarity.**

Beta dispersion significance for the Bray-Curtis distances of the developmental comparison of pup (large intestinal wash) samples. *P. polionotus* and *P. maniculatus* samples differ in dispersion by timepoint for community structure.

| Subset | | Df | SumofSqs | Mean Sq | F value | N.Perm | Pr(>F) |
|---|---|---|---|---|---|---|---|
| CA | Timepoint | 2 | 0.075 | 0.038 | 1.47 | 99 | 0.18 |
| | Residuals | 15 | 0.38 | 0.025 | | | |
| LE | Timepoint | 2 | 0.058 | 0.029 | 1.65 | 99 | 0.25 |
| | Residuals | 17 | 0.30 | 0.018 | | | |
| PO | Timepoint | 2 | 0.20 | 0.099 | 9.28 | 99 | **0.01** |
| | Residuals | 14 | 0.15 | 0.011 | | | |
| MA | Timepoint | 2 | 0.18 | 0.089 | 18.4 | 99 | **0.01** |
| | Residuals | 18 | 0.087 | 0.0048 | | | |

**Table P. Kruskal-Wallis tests of pairwise distances between pup and adult samples, Jaccard distances.**

| Subset | Pup group distance to: | Kruskal-Wallis P | Number of Comparisons | Number of Values | Kruskal-Wallis Statistic |
|---|---|---|---|---|---|
| CA | Adult Fecal | **< 0.0001** | 3 | 108 | 80.3 |
| | Adult Mammary Skin | **< 0.0001** | 3 | 108 | 32.7 |
| LE | Adult Fecal | **< 0.0001** | 3 | 100 | 23.7 |
| | Adult Mammary Skin | **0.0025** | 3 | 240 | 12.0 |
| PO | Adult Fecal | **< 0.0001** | 3 | 85 | 71.5 |
| | Adult Mammary Skin | **< 0.0001** | 3 | 102 | 54.5 |
| MA | Adult Fecal | **< 0.0001** | 3 | 168 | 58.0 |
| | Adult Mammary Skin | **< 0.0001** | 3 | 147 | 19.0 |

**Table Q. Dunn's multiple comparisons: Pup sample Jaccard distances to parental fecal samples.**

| | | CA | | LE | | PO | | MA | |
|---|---|---|---|---|---|---|---|---|---|
| Group 1 | Group 2 | Mean rank diff. | Adj. P Value | Mean rank diff. | Adj. P Value | Mean rank diff. | Adj. P Value | Mean rank diff. | Adj. P Value |
| Newborn | Pre-weaning | 31.7 | **< 0.0001** | 18.9 | **0.0382** | 26.6 | **0.0002** | 29.1 | **0.0053** |
| Newborn | Weaning | 66.1 | **< 0.0001** | 35.8 | **< 0.0001** | 53.9 | **< 0.0001** | 67.7 | **< 0.0001** |
| Pre-weaning | Weaning | 34.4 | **< 0.0001** | 16.9 | **0.0361** | 27.3 | **0.0001** | 38.7 | **0.0002** |

**Table R. Dunn's multiple comparisons: Pup sample Jaccard distances to maternal mammary skin samples.**

| | | CA | | LE | | PO | | MA | |
|---|---|---|---|---|---|---|---|---|---|
| Group 1 | Group 2 | Mean rank diff. | Adj. P Value | Mean rank diff. | Adj. P Value | Mean rank diff. | Adj. P Value | Mean rank diff. | Adj. P Value |
| Newborn | Pre-weaning | 31.5 | **< 0.0001** | -10.5 | > 0.99 | 26.3 | **0.0010** | 20.4 | **0.0569** |
| Newborn | Weaning | 40.1 | **< 0.0001** | 24.5 | 0.0956 | 51.5 | **< 0.0001** | 36.1 | **< 0.0001** |
| Pre-weaning | Weaning | 8.64 | 0.7257 | 35.0 | **0.0022** | 25.2 | **0.0017** | 15.7 | 0.2386 |

**Table S. Kruskal-Wallis tests of pairwise distances between pup and adult samples, Bray-Curtis distances.**

| Subset | Pup group distance to: | Kruskal-Wallis P | Number of Comparisons | Number of Values | Kruskal-Wallis Statistic |
|---|---|---|---|---|---|
| CA | Adult Fecal | **< 0.0001** | 3 | 117 | 96.4 |
| | Adult Mammary Skin | **< 0.0001** | 3 | 108 | 28.1 |
| LE | Adult Fecal | 0.535 | 3 | 100 | 1.25 |
| | Adult Mammary Skin | **0.0215** | 3 | 120 | 7.68 |
| PO | Adult Fecal | **< 0.0001** | 3 | 85 | 57.3 |
| | Adult Mammary Skin | **< 0.0001** | 3 | 102 | 42.7 |
| MA | Adult Fecal | **< 0.0001** | 3 | 168 | 30.3 |
| | Adult Mammary Skin | **0.0007** | 3 | 147 | 14.5 |

**Table T. Dunn's multiple comparisons: Pup sample Bray-Curtis distances to parental fecal samples.**

| | | CA | | LE | | PO | | MA | |
|---|---|---|---|---|---|---|---|---|---|
| Group 1 | Group 2 | Mean rank diff. | Adj. P Value | Mean rank diff. | Adj. P Value | Mean rank diff. | Adj. P Value | Mean rank diff. | Adj. P Value |
| Newborn | Pre-weaning | 39.9 | **< 0.0001** | 8.50 | 0.7900 | 27.5 | **0.0001** | 24.3 | **0.0268** |
| Newborn | Weaning | 75.4 | **< 0.0001** | 4.92 | > 0.99 | 48.1 | **< 0.0001** | 49.0 | **< 0.0001** |
| Pre-weaning | Weaning | 35.5 | **< 0.0001** | -3.58 | > 0.99 | 20.5 | **0.0064** | 24.7 | **0.0293** |

**Table U. Dunn's multiple comparisons: Pup sample Bray-Curtis distances to maternal mammary skin samples.**

| | | CA | | LE | | PO | | MA | |
|---|---|---|---|---|---|---|---|---|---|
| Group 1 | Group 2 | Mean rank diff. | Adj. P Value | Mean rank diff. | Adj. P Value | Mean rank diff. | Adj. P Value | Mean rank diff. | Adj. P Value |
| Newborn | Pre-weaning | -0.0278 | > 0.99 | -10.5 | > 0.99 | 24.7 | **0.0022** | 9.59 | 0.809 |
| Newborn | Weaning | 33.9 | **< 0.0001** | 24.5 | 0.0956 | 45.5 | **< 0.0001** | 31.2 | **0.0005** |
| Pre-weaning | Weaning | 33.9 | **< 0.0001** | 35.0 | **0.0022** | 20.8 | **0.0134** | 21.6 | **0.0469** |

**Table V. Kruskal-Wallis tests of pairwise pup samples, Jaccard distances.**

| Subset | Pup distance to: | Kruskal-Wallis P | Number of Comparisons | Number of Values | Kruskal-Wallis Statistic |
|---|---|---|---|---|---|
| CA | other pup timepoints | **< 0.0001** | 3 | 108 | 40.9 |
| LE | other pup timepoints | **0.0020** | 3 | 131 | 12.4 |
| PO | other pup timepoints | **< 0.0001** | 3 | 96 | 42.0 |
| MA | other pup timepoints | **0.0002** | 3 | 146 | 16.8 |

**Table W. Kruskal-Wallis tests of pairwise pup samples, Bray-Curtis distances.**

| Subset | Pup distance to: | Kruskal-Wallis P | Number of Comparisons | Number of Values | Kruskal-Wallis Statistic |
|---|---|---|---|---|---|
| CA | other pup timepoints | **< 0.0001** | 3 | 108 | 41.5 |
| LE | other pup timepoints | **< 0.0001** | 3 | 131 | 21.4 |
| PO | other pup timepoints | **< 0.0001** | 3 | 96 | 34.4 |
| MA | other pup timepoints | **0.0004** | 3 | 146 | 15.7 |

**Table X. Dunn's multiple comparisons: Pup sample Jaccard distances between timepoints.**

| | | CA | | LE | |
|---|---|---|---|---|---|
| **Comparison 1** | **Comparison 2** | Mean rank diff. | Adj. P Value | Mean rank diff. | Adj. P Value |
| Newborn x Pre-weaning | Newborn x Weaning | -36.7 | **< 0.0001** | -28.4 | **0.0037** |
| Newborn x Pre-weaning | Pre-weaning x Weaning | 7.42 | 0.945 | -5.78 | > 0.99 |
| Newborn x Weaning | Pre-weaning x Weaning | 44.1 | **< 0.0001** | 22.6 | **0.0119** |

**Table Y. Dunn's multiple comparisons: Pup sample Jaccard distances between timepoints (continued).**

| | | PO | | MA | |
|---|---|---|---|---|---|
| **Comparison 1** | **Comparison 2** | Mean rank diff. | Adj. P Value | Mean rank diff. | Adj. P Value |
| Newborn x Pre-weaning | Newborn x Weaning | -33.7 | **< 0.0001** | -30.2 | **0.0008** |
| Newborn x Pre-weaning | Pre-weaning x Weaning | 7.75 | 0.844 | -1.67 | > 0.99 |
| Newborn x Weaning | Pre-weaning x Weaning | 41.4 | **< 0.0001** | 28.6 | **0.0028** |

**Table Z. Dunn's multiple comparisons: Pup sample Bray-Curtis distances between timepoints.**

| | | CA | | LE | |
|---|---|---|---|---|---|
| **Comparison 1** | **Comparison 2** | Mean rank diff. | Adj. P Value | Mean rank diff. | Adj. P Value |
| Newborn x Pre-weaning | Newborn x Weaning | -44.9 | **< 0.0001** | -31.8 | **0.0009** |
| Newborn x Pre-weaning | Pre-weaning x Weaning | -8.81 | 0.699 | -36.3 | **< 0.0001** |
| Newborn x Weaning | Pre-weaning x Weaning | 36.1 | **< 0.0001** | -4.42 | > 0.99 |

**Table AA. Dunn's multiple comparisons: Pup sample Bray-Curtis distances between timepoints (continued).**

| | | PO | | MA | |
|---|---|---|---|---|---|
| **Comparison 1** | **Comparison 2** | Mean rank diff. | Adj. P Value | Mean rank diff. | Adj. P Value |
| Newborn x Pre-weaning | Newborn x Weaning | -14.7 | 0.0968 | -32.1 | **0.0003** |
| Newborn x Pre-weaning | Pre-weaning x Weaning | 25.4 | **0.0012** | -24.2 | **0.0205** |
| Newborn x Weaning | Pre-weaning x Weaning | 40.2 | **< 0.0001** | 7.93 | > 0.99 |

#### Appendix C4. Host transcriptomic analyses – Full sample set.

**Table BB. ADONIS results for full host transcriptomics sample set.**

ADONIS statistics on euclidean distances wherein species is *CA*, *LE*, *PO*, and *MA*, and timepoint newborn, pre-weaning, weaning, or post-weaning. Adonis2 formula of Jaccard ~ Species + SampleType, with 1 x $10^5$ permutations and "by = margin". Both host species and timepoint explain a significant amount of the beta diversity in the sample set.

| | | Df | SumofSqs | $R^2$ | F | Pr(>F) |
|---|---|---|---|---|---|---|
| ***PCA vst*** | Species | 3 | 20479 | 0.483 | 123 | **< 0.0001** |
| | Timepoint | 2 | 18078 | 0.426 | 162 | **< 0.0001** |
| | Residual | 50 | 2786 | 0.0657 | | |

#### Appendix C5. Host transcriptomic analyses – Developmental comparison.

**Table CC. ADONIS results for euclidean distances.**

ADONIS statistics on euclidean distances principal components for the developmental comparison of pup (large intestinal tissue) samples. Adonis2 formula of PCA ~ Timepoint, with 1 x $10^5$ permutations and "by = margin". Developmental timepoint explains a significant amount of beta diversity variation of host gene expression profiles for all *Peromyscus* species.

| Subset | | Df | SumofSqs | $R^2$ | F | Pr(>F) |
|---|---|---|---|---|---|---|
| *CA* | Timepoint | 2 | 7041 | 0.710 | 9.78 | **0.00076** |
| | Residual | 8 | 2879 | 0.290 | | |
| *LE* | Timepoint | 2 | 10154 | 0.896 | 60.5 | **< 0.0001** |
| | Residual | 14 | 1174 | 0.104 | | |
| *PO* | Timepoint | 2 | 9622 | 0.736 | 13.9 | **< 0.0001** |
| | Residual | 10 | 3453 | 0.264 | | |
| *MA* | Timepoint | 2 | 12860 | 0.765 | 19.6 | **< 0.0001** |
| | Residual | 12 | 3944 | 0.235 | | |

**Table DD. Average distance to centroid for euclidean distances.**

Beta dispersion estimates on euclidean distances for the developmental comparison of pup (large intestinal tissue) samples. Average distance to the centroid for each species and timepoint is reported.

| Large Intestine Tissue | Newborn Pups | Pre-weaning Pups | Weaning Pups |
|---|---|---|---|
| CA | 14.8 | 10.7 | 19.1 |
| LE | 7.75 | 9.05 | 5.84 |
| PO | 19.9 | 10.9 | 6.21 |
| MA | 16.1 | 10.8 | 7.32 |

**Table EE. Permutation test for homogeneity of multivariate dispersions for euclidean distances.**

Beta dispersion significance for the euclidean distances of the developmental comparison of pup (large intestinal tissue) samples. Only *P. polionotus* differs in dispersion by timepoint.

| Subset | | Df | SumofSqs | Mean Sq | F value | N.Perm | Pr(>F) |
|---|---|---|---|---|---|---|---|
| CA | Timepoint | 2 | 99 | 49 | 0.663 | 99 | 0.54 |
| | Residuals | 8 | 597 | 75 | | | |
| LE | Timepoint | 2 | 33 | 17 | 1.49 | 99 | 0.25 |
| | Residuals | 14 | 156 | 11 | | | |
| PO | Timepoint | 2 | 431 | 216 | 4.50 | 99 | **0.04** |
| | Residuals | 10 | 479 | 48 | | | |
| MA | Timepoint | 2 | 214 | 107 | 0.778 | 99 | 0.47 |
| | Residuals | 12 | 1654 | 138 | | | |

#### Appendix C6. Microbial beta diversity – Host species comparison.

**Table FF. ADONIS results for Jaccard distances.**

ADONIS statistics on Jaccard distances for the host species comparison of pup (large intestinal wash) samples. Adonis2 formula of Jaccard ~ Species, with 1 x $10^5$ permutations and “by = margin”.

| *Subset* | | Df | SumofSqs | $R^2$ | F | Pr(>F) |
|---|---|---|---|---|---|---|
| *Newborn* | Species | 3 | 1.7 | 0.167 | 1.40 | **0.0036** |
| | Residual | 21 | 8.5 | 0.833 | | |
| *Pre-weaning* | Species | 3 | 1.7 | 0.196 | 1.63 | **< 0.0001** |
| | Residual | 20 | 6.8 | 0.804 | | |
| *Weaning* | Species | 3 | 2.0 | 0.204 | 1.96 | **0.00027** |
| | Residual | 23 | 7.8 | 0.796 | | |

**Table GG. Average distance to centroid for Jaccard similarity.**

Beta dispersion estimates on Jaccard distances for the host species comparison of pup (large intestinal wash) samples. Average distance to the centroid for each timepoint and species is reported.

| Large Intestine Wash | CA | LE | PO | MA |
|---|---|---|---|---|
| Newborn | 0.547 | 0.541 | 0.597 | 0.617 |
| Pre-weaning | 0.532 | 0.577 | 0.509 | 0.492 |
| Weaning | 0.491 | 0.560 | 0.510 | 0.561 |

**Table HH. Permutation test for homogeneity of multivariate dispersions for Jaccard similarity.**

Beta dispersion significance for the Jaccard distances of the host species comparison of pup (large intestinal wash). Only weaning pups differ significantly in beta dispersion by host species for community membership.

| Subset | | Df | SumofSqs | Mean Sq | F value | N.Perm | Pr(>F) |
|---|---|---|---|---|---|---|---|
| Newborn | Groups | 3 | 0.026 | 0.0088 | 2.10 | 99 | 0.08 |
| | Residuals | 21 | 0.088 | 0.0042 | | | |
| Pre-weaning | Groups | 3 | 0.026 | 0.0087 | 3.03 | 99 | 0.09 |
| | Residuals | 20 | 0.058 | 0.0029 | | | |
| Weaning | Groups | 3 | 0.025 | 0.0084 | 3.73 | 99 | **0.04** |
| | Residuals | 23 | 0.052 | 0.0023 | | | |

**Table II. ADONIS results for Bray-Curtis distances.**

ADONIS statistics on Bray-Curtis distances for the host species comparison of pup (large intestinal wash) samples. Adonis2 formula of Bray ~ Species, with 1 x $10^5$ permutations and "by = margin".

| Subset | | Df | SumofSqs | $R^2$ | F | Pr(>F) |
|---|---|---|---|---|---|---|
| Newborn | Species | 3 | 2.0 | 0.252 | 2.35 | **0.0069** |
| | Residual | 21 | 5.8 | 0.748 | | |
| Pre-weaning | Species | 3 | 1.3 | 0.256 | 2.30 | **0.0003** |
| | Residual | 20 | 3.8 | 0.744 | | |
| Weaning | Species | 3 | 2.3 | 0.282 | 3.01 | **< 0.0001** |
| | Residual | 23 | 5.8 | 0.718 | | |

**Table JJ. Average distance to centroid for Bray-Curtis dissimilarity.**

Beta dispersion estimates on Bray-Curtis distances for the host species comparison of pup (large intestinal wash) samples. Average distance to the centroid for each timepoint and species is reported.

| | Large Intestine Wash | | | |
|---|---|---|---|---|
| | CA | LE | PO | MA |
| Newborn | 0.300 | 0.370 | 0.576 | 0.512 |
| Pre-weaning | 0.415 | 0.458 | 0.305 | 0.292 |
| Weaning | 0.443 | 0.504 | 0.409 | 0.457 |

**Table KK. Permutation test for homogeneity of multivariate dispersions for Bray-Curtis dissimilarity.**

Beta dispersion significance for the Bray-Curtis distances of the host species comparison of pup (large intestinal wash). Only newborn and pre-weaning pups differ in beta dispersion by host species for community structure.

| Subset | | Df | SumofSqs | Mean Sq | F value | N.Perm | Pr(>F) |
|---|---|---|---|---|---|---|---|
| Newborn | Groups | 3 | 0.29 | 0.097 | 4.10 | 99 | **0.01** |
| | Residuals | 21 | 0.50 | 0.024 | | | |
| Pre-weaning | Groups | 3 | 0.12 | 0.041 | 2.76 | 99 | **0.04** |
| | Residuals | 20 | 0.30 | 0.015 | | | |
| Weaning | Groups | 3 | 0.033 | 0.011 | 2.33 | 99 | 0.07 |
| | Residuals | 23 | 0.11 | 0.0047 | | | |

#### Appendix C7. Host transcriptomic beta diversity – Host species comparison.

**Table LL. ADONIS results for euclidean distances.**

ADONIS statistics on euclidean distances principal components for the host species comparison of pup (large intestinal tissue) samples. Adonis2 formula of PCA ~ Species, with 1 x $10^5$ permutations and "by = margin".

| Subset | | Df | SumofSqs | $R^2$ | F | Pr(>F) |
|---|---|---|---|---|---|---|
| Newborn | Species | 3 | 11804 | 0.651 | 10.6 | **< 0.0001** |
| | Residual | 17 | 6323 | 0.349 | | |
| Pre-weaning | Species | 3 | 15104 | 0.968 | 149 | **< 0.0001** |
| | Residual | 15 | 507 | 0.0325 | | |
| Weaning | Species | 3 | 10013 | 0.969 | 125 | **< 0.0001** |
| | Residual | 12 | 320 | 0.0310 | | |

**Table MM. Average distance to centroid for euclidean distances.**

Beta dispersion estimates on euclidean distances for the host species comparison of pup (large intestinal tissue) samples. Average distance to the centroid for each timepoint and species is reported.

| | Large Intestinal Tissue | | | |
|---|---|---|---|---|
| | CA | LE | PO | MA |
| Newborn | 9.0 | 9.1 | 17.1 | 14.8 |
| Pre-weaning | 5.6 | 4.5 | 2.1 | 5.9 |
| Weaning | 6.2 | 2.9 | 3.4 | 4.3 |

**Table NN. Permutation test for homogeneity of multivariate dispersions for euclidean distances.**

Beta dispersion significance for euclidean distances of the host species comparison of pup (large intestinal tissue) samples. No timepoints differ in beta dispersion across host species.

| Subset | | Df | SumofSqs | Mean Sq | F value | N.Perm | Pr(>F) |
|---|---|---|---|---|---|---|---|
| Newborn | Groups | 3 | 258 | 86 | 0.579 | 99 | 0.69 |
| | Residuals | 17 | 2529 | 149 | | | |
| Pre-weaning | Groups | 3 | 36 | 12 | 2.15 | 99 | 0.15 |
| | Residuals | 15 | 84 | 5.6 | | | |
| Weaning | Groups | 3 | 17 | 5.8 | 1.06 | 99 | 0.52 |
| | Residuals | 12 | 65 | 5.4 | | | |

**Table OO. Comparison of expression divergence rates of newborn gene modules.**

For each module (Orchid, Skyblue, and Honeydew), analysis of covariance (ANCOVA) was conducted using the formula (expr_distance ~ phylo_distance * module), slopes compared using two-tailed t-tests, and multiple comparisons corrected for using Benjamini-Hochberg.

| | Unweighted Expression | | | Weighted Expression | | |
|---|---|---|---|---|---|---|
| **Modules** | Slope | Difference | Adj. P | Slope | Difference | Adj. P |
| Honeydew<br>Orchid | 1.027<br>0.332 | -0.70 | **0.012** | 36.696<br>5.181 | -31.51 | **0.00008** |
| Honeydew<br>Skyblue | 1.027<br>1.117 | 0.090 | 0.69 | 36.696<br>21.916 | -14.78 | **0.0097** |
| Orchid<br>Skyblue | 0.332<br>1.117 | 0.79 | **0.011** | 5.181<br>21.916 | 16.74 | **0.0069** |

**Table PP. Jonckheere-terpstra test of monotonic trend on alpha diversity.**

For each species and alpha statistic, a Jonckheere-terpstra test was conducted using 'alternative = "increasing"', and 'nperm = 5000'. Each alpha statistic was tested to determine if there was an increasing monotonic trend across pup age ("timepoint").

| Alpha Statistic | Subset | JT | P-value |
|---|---|---|---|
| Unique ASVs | CA | 99 | **0.0002** |
| | LE | 97.5 | **0.0126** |
| | PO | 76.5 | **0.0046** |
| | MA | 79.5 | 0.3452 |
| Shannon Diversity | CA | 104 | **0.0002** |
| | LE | 107 | **0.0012** |
| | PO | 88 | **0.0002** |
| | MA | 94 | 0.0906 |
| Faiths PD | CA | 97 | **0.0002** |
| | LE | 95 | **0.0226** |
| | PO | 77 | **0.0048** |
| | MA | 69 | 0.6174 |
| Pielou's Evenness | CA | 102 | **0.0002** |
| | LE | 103 | **0.0038** |
| | PO | 90 | **0.0004** |
| | MA | 102 | **0.032** |
